# A cooperative network at the nuclear envelope counteracts LINC-mediated forces during oogenesis in *C. elegans*

**DOI:** 10.1101/2021.08.13.456272

**Authors:** Chenshu Liu, Zoe Lung, John S. Wang, Fan Wu, Abby F. Dernburg

**Author notes:** ABBREVIATIONS: NE – Nuclear Envelope; LINC – LInker of Nucleoskeleton and Cytoskeleton; MT – microtubules; INM – inner nuclear membrane; ONM – outer nuclear membrane; SC – synaptonemal complex; TZ – transition zone; EP – early pachytene; LP – late pachytene; Dip – diplotene; AID – auxin-inducible degradation; LEM – LAP2, emerin, MAN1.

## Abstract

Oogenesis involves meiosis and oocyte maturation. Both processes rely on mechanical forces (Lee et al., 2015; Nagamatsu et al., 2019; Rog and Dernburg, 2015; Sato et al., 2009; Tsatskis et al., 2020; Wynne et al., 2012), which can be transduced from the cytoskeleton to the nuclear envelope (NE) through linker of nucleoskeleton and cytoskeleton (LINC) complexes (Burke, 2018; Chang et al., 2015; Fan et al., 2020; Link et al., 2014). Gametes must protect their genomes from damage in this mechanically stressful environment. In *C. elegans,* oocyte nuclei lacking the single lamin protein LMN-1 are vulnerable to nuclear collapse. Here we deploy the auxin-inducible degradation system to investigate the balance of forces that drive this collapse and protect oocyte nuclei. We find that nuclear collapse is not a consequence of apoptosis. It is promoted by dynein and a LINC complex comprised of SUN-1 and ZYG-12, which assumes polarized distribution at the NE in response to dynein-mediated forces. We also show that the lamin meshwork works in parallel with other inner nuclear membrane (INM) proteins to counteract mechanical stress at the NE during oogenesis. We speculate that a similar network may protect oocyte integrity during the long arrest period in mammals.

## INTRODUCTION

Oogenesis produces haploid ova from diploid progenitors through meiosis and oocyte differentiation and maturation. During meiosis, homologous chromosomes must pair, synapse and undergo controlled DNA double strand breaks (DSBs) and repair (Bhalla and Dernburg, 2008; Yu et al., 2016). Oocyte nuclei are subjected to mechanical forces generated by the cytoskeleton and transmitted to the nuclear envelope through LINC complexes (Lee et al., 2015; Link et al., 2014; Luo et al., 2016; Sato et al., 2009; Wynne et al., 2012). These forces are important for meiotic chromosome dynamics and for the positioning and movement of nuclei and oocytes in the gonad during maturation(Zhou et al., 2009). Mammalian oocytes undergo homolog pairing, synapsis, and recombination in embryonic ovaries and then arrest at the diplotene stage of meiotic prophase. Following puberty, hormonal cycles lead to oocyte growth and ovulation. This arrest can persist for decades in humans and other mammals with long reproductive lifespans (Zhang, 2018). Mechanical compression has been shown to play a key role in maintaining this dormant state in mice (Nagamatsu et al., 2019). During the long diplotene arrest, loss of NE stiffness correlates with diminished reserve of dormant oocytes(Tsatskis et al., 2020). However, the molecular mechanisms that confer NE stiffness in diplotene oocytes, and the fate of oocyte nuclei when NE stiffness is compromised are not well-studied.

The nuclear lamina is a meshwork comprised of lamins, type V intermediate filament proteins (Aebi et al., 1986; Goldman et al., 1986; McKeon et al., 1986). Lamins self-assemble into higher-order structures which confer mechanical stability to the NE and protect nuclear contents (Davidson and Lammerding, 2014; Swift et al., 2013). Lamin mutations cause diverse diseases collectively called laminopathies (Broers et al., 2006; Worman, 2012). Defects in the nuclear lamina can cause nuclear rupture, DNA damage, alterations in chromosomal architecture or loss of genome integrity in somatic tissues (Denais et al., 2016; Earle et al., 2020; Hatch et al., 2013; Kneissig et al., 2019).

The lamin meshwork is present in metazoan oocytes and likely reinforces the NE during oogenesis (Link et al., 2013; Link et al., 2018). Consistent with this idea, women carrying *LMNA* mutations are at higher risks for infertility and obstetric complications (Vantyghem et al., 2008). In mammals, the roles of nuclear lamina during oogenesis is confounded by the presence of multiple lamin isoforms in the germline (Link et al., 2013). *C. elegans* has only one lamin protein (LMN-1). Null mutations in *lmn-1* result in near-complete sterility and embryonic lethality, with hypercondensed chromatin observed in oocyte nuclei (Green et al., 2011; Huelgas-Morales et al., 2020; Liu et al., 2000). However, pleiotropic effects on both somatic and germline tissues in the *lmn-1* null mutant, such as atrophic gonads and accelerated aging, have obscured the direct roles of LMN-1 during oogenesis. To examine the germline specific functions of LMN-1, we used the auxin-inducible degradation system to enable conditional and tissue specific depletion of LMN-1(Zhang et al., 2015).

## RESULTS

### Induced degradation of LMN-1 in the *C. elegans* germline causes apoptosis-independent nuclear collapse in maturing oocytes

Lamin protein-protein interactions are easily perturbed by the addition of epitope tags or other sequences (Velez-Aguilera et al., 2020). However, we inserted a minimal auxin-inducible degron (AID) with a V5 epitope tag into a poorly conserved yet highly flexible linker region within LMN-1 (Dechat et al., 2010) in a strain expressing TIR1 in the germline, and found that homozygous animals were fully viable and fertile in the absence of auxin (**Figure 1a; Supplementary Figure 1a; Supplementary Table 1**). When exposed to indole acetic acid (IAA, auxin), we observed robust depletion of LMN-1 in the germline; the signal remaining in the somatic distal tip cell and germline sheath cells also provided a convenient positive control for immunofluorescence. This resulted in complete inviability of embryos laid on auxin-containing plates (**Figure 1b,c; Supplementary Figure 1b**).

**Figure 1:**
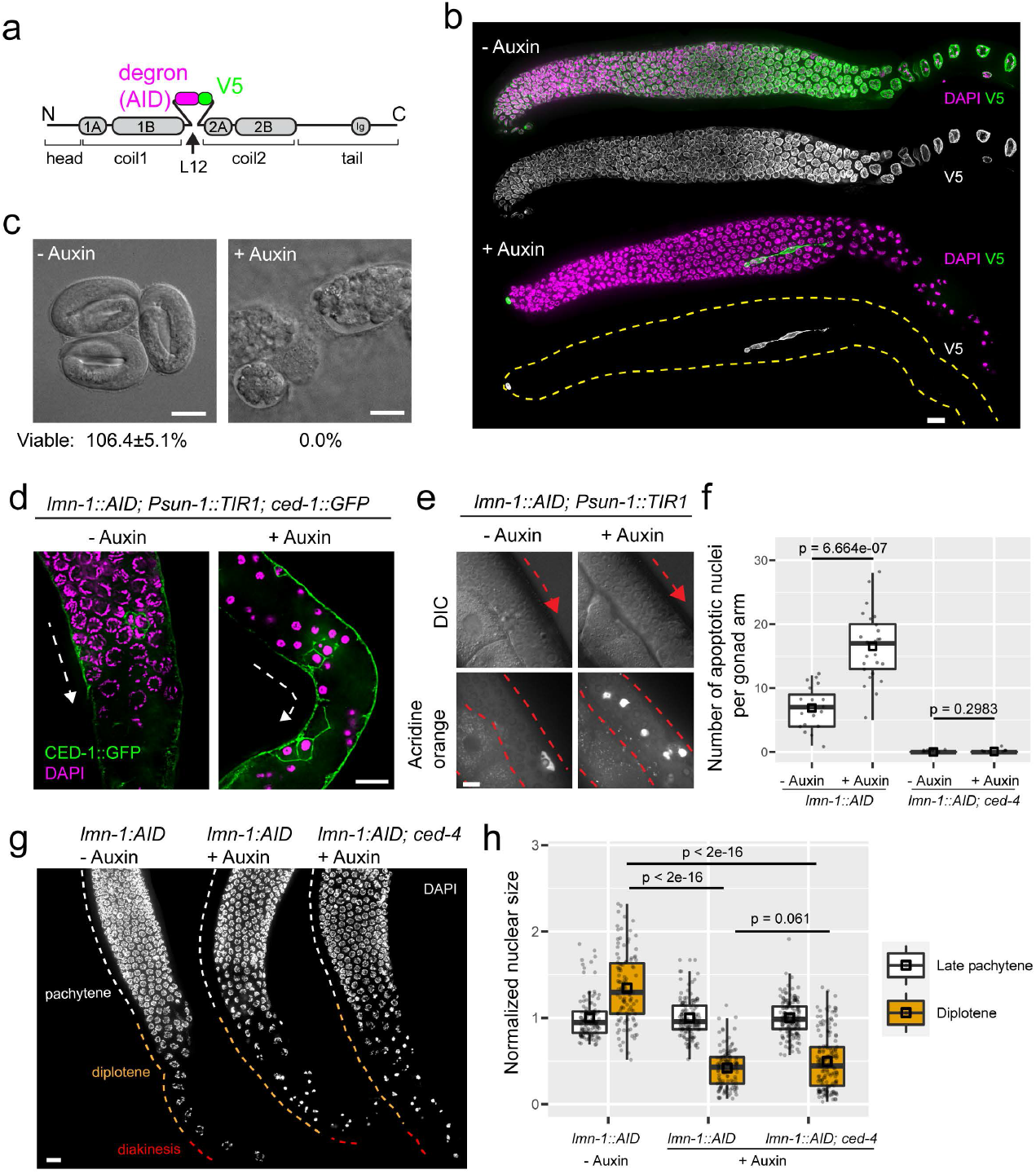
Induced degradation of LMN-1 in the *C. elegans* germline causes apoptosis-independent nuclear collapse in maturing oocytes. **a.** Domain organization of *C. elegans* LMN-1, indicating the position where a degron (AID) and V5-epitope were inserted. **b.** LMN-1::V5::AID immunofluorescence in gonads dissected from adult hermaphrodites. Meiosis progresses from left to right. DAPI, magenta; anti-V5, green. Grayscale images show LMN-1::V5::AID alone. Following auxin treatment, LMN-1::V5::AID is detected only in the distal tip cell and sheath cells. Images are maximum-intensity projections and scaled identically. Scale bar, 10 µm. **c.** Differential interference contrast (DIC) images of embryos laid by hermaphrodites following auxin treatment for 48 hours, and (–)auxin controls. Scale bar, 20 µm. **d.** Immunofluorescence of CED-1::GFP. Following depletion of LMN-1 depletion, nuclei in the proximal gonad have hypercondensed chromatin, and a subset are surrounded by CED-1::GFP, a marker for engulfing cells. Dashed lines indicate the direction of meiotic progression. Scale bars, 10 µm. **e.** Images of germlines in living animals showing gonad morphology (DIC) and corresponding apoptotic nuclei stained with acridine orange. Red dashed lines and arrows indicate the contour of gonad and the direction of meiotic progression. Scale bar, 10 µm. **f.** Quantification of apoptotic nuclei per gonad arm. All worms were homozygous for *P_sun-1_::TIR1*. Each dot represents one animal. Medians (black crossbars) and means (black boxes) are shown. *p*-values were calculated using the Mann-Whitney test. **g.** Nuclear morphology during late meiotic prophase. All worms were homozygous for *P_sun-1_::TIR1*. Meiotic prophase progresses from top to bottom; stages are annotated based on their anatomical positions within control gonads. Scale bar, 10 µm. **h.** Quantification of nuclear size. Medians (black crossbars) and means (black boxes) are shown. *p*-values were calculated using one-way ANOVA and *post hoc* pairwise *t*-tests (two-sided; adjusted by the Benjamini-Hochberg method). Colors correspond to the regions shown in (**g**).

Depletion of LMN-1 recapitulated germline defects previously described in *lmn-1* null mutants: dramatic compaction of nuclei in the proximal region near the uterus, in the region corresponding to diplotene and diakinesis in wild-type hermaphrodites (Green et al., 2011; Huelgas-Morales et al., 2020; Liu et al., 2000). This nuclear collapse is characterized by a drastic reduction in chromatin volume, which normally increases gradually from late pachytene to diplotene (**Figure 1b,g,h; Supplementary Figures 2,3**). We also observed persistent RAD-51 foci, a marker for DNA double strand break (DSB) repair intermediates (Rinaldo et al., 2002). While the majority of these RAD-51 foci depended on the meiosis specific endonuclease SPO-11, both RAD-51 and GFP::COSA-1 foci, which mark designated crossover sites (Yokoo et al., 2012), were also detected in the absence of SPO-11 (**Supplementary Figure 4**). This is consistent with previous evidence that disruption of the nuclear lamina can cause DNA damage (Denais et al., 2016; Earle et al., 2020).

Because unrepaired DNA damage in oocyte nuclei causes increased apoptosis in *C. elegans* germline (Bohr et al., 2016; Penkner et al., 2007), and apoptotic nuclei have hypercondensed chromatin morphology (Raiders et al., 2018), we investigated whether the condensed nuclei observed following LMN-1 depletion were apoptotic using two different markers. We stained live worms with acridine orange (Penkner et al., 2007), and also crossed in a fluorescent CED-1::GFP marker, which localizes to the membranes of apoptotic cells undergoing engulfment (Bhalla and Dernburg, 2005). Both markers were sharply increased in hermaphrodite germlines depleted of LMN-1 (**Figure 1d-f**). However, we observed condensation of nuclear DNA in this region even in the absence of CED-4, which is essential for germline apoptosis (**Figure 1f-h**) (Bhalla and Dernburg, 2005). Together with studies of several known meiotic mutants that result in elevated apoptosis but not collapsed nuclei in diplotene (**Supplementary Figure 5**) (Bhalla and Dernburg, 2005; Bohr et al., 2016; Bohr et al., 2015; Gartner et al., 2000; MacQueen et al., 2002), our results demonstrate that nuclear collapse is not a consequence of apoptosis.

### Nuclear collapse is driven by force-induced destabilization of the nuclear envelope

To better understand the process leading to nuclear collapse, we introduced fluorescent nuclear envelope markers SUN-1::mRuby or ZYG-12::GFP to enable live-cell imaging following LMN-1 depletion (**Supplementary Video 1)**. We also combined ZYG-12::GFP with a fluorescently labeled synaptonemal complex (SC) protein, mRuby::SYP-3, to enable observation of chromosomes (**Figure 2**) (Rog et al., 2017). The LINC complex components SUN-1 and its KASH partner ZYG-12 span the inner nuclear membrane (INM) and outer nuclear membrane (ONM) respectively and play important roles in connecting chromosomes to the microtubule cytoskeleton during early meiosis in the distal region of the germline (Sato et al., 2009; Wynne et al., 2012). During early prophase, SUN-1 and ZYG-12 connect specialized chromosome regions known as meiotic pairing centers to cytoplasmic dynein to promote movement of chromosomes along the nuclear envelope. This movement is important for homolog pairing and synapsis, and normally abates once chromosomes are fully synapsed (Wynne et al., 2012). In germlines depleted of LMN-1, although there was no delay in pairing or synapsis (**Supplementary Figure 6**), clustered LINC complex persisted from early meiosis through the proximal region (**Figure 2a-c; Supplementary Figure 7**). Particle tracking in live animals revealed that these patches remained highly mobile within the nuclear envelope throughout meiotic prophase (**Figure 2d,e; Supplementary Videos 2,3**). LMN-1 is normally phosphorylated during early meiosis and then dephosphorylated upon synapsis as chromosome movements subside(Link et al., 2018); our observations indicate that the forces transmitted to the NE do not cease at this time but that the lamina restricts movement of LINC complexes and pairing centers as dephosphorylation leads to more extensive crosslinking.

**Figure 2:**
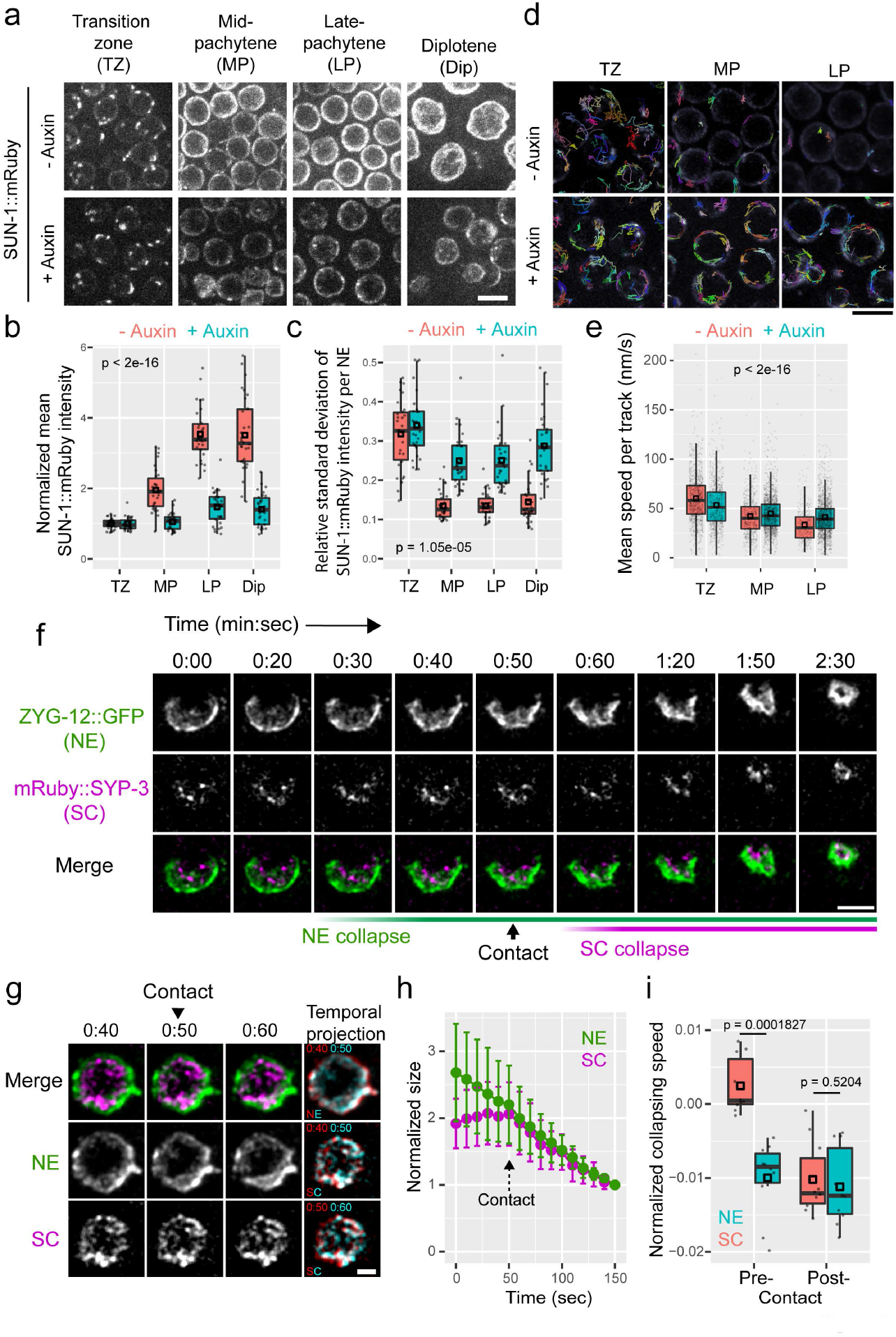
Dynamics of nuclear collapse. **a-e.** Prolonged LINC complex clustering and mobility following LMN-1 depletion**. a**, Maximum intensity projection images of SUN-1::mRuby at different stages of meiotic prophase. Images are scaled identically. Scale bar, 5 µm. **b.** Mean intensities of SUN-1::mRuby at the nuclear envelope per nucleus, normalized against intensity levels at the transition zone. Medians (black crossbars) and means (black boxes) are shown. Fluorescence intensity surrounding each nucleus was manually segmented and quantified from additive projection images after background subtraction. Three animals, each with 10 meiotic nuclei per stage, were measured per condition (without or with auxin). TZ, transition zone; MP, mid-pachytene; LP, late pachytene; Dip, diplotene. Two-way ANOVA was used to calculate the *p*-value. **c.** Clustering of SUN-1::mRuby, defined as the relative standard deviation (ratio of the standard deviation to the mean value) of fluorescence intensity at the NE in each nucleus. Medians (black crossbars) and means (black boxes) are shown. Fluorescence intensity was measured in the same way as in (**b**). Three animals, each with 10 meiotic nuclei per stage, were measured per condition. TZ, transition zone; MP, mid-pachytene; LP, late pachytene; Dip, diplotene. *<u>p</u>*-value was calculated by two-way ANOVA. **d.** Grayscale images of SUN-1::mRuby overlaid with color-coded trajectories generated by 3D-particle-tracking of SUN-1::mRuby patches. Scale bar, 5 µm. **e.** Mobility of SUN-1::mRuby patches at the NE, quantified as the mean speed of individual patches. Medians (black crossbars) and means (black boxes) are shown. *p*-value was calculated with two-way ANOVA. Three animals were measured per condition per stage. TZ, transition zone (n = 1132 detected particles for ‘- Auxin’, n = 859 for ‘+ Auxin’); MP, mid-pachytene (n = 620 detected particles for ‘- Auxin’, n = 1657 for ‘+ Auxin’); LP, late pachytene (n = 306 detected particles for ‘- Auxin’, n = 1675 for ‘+ Auxin’). **f.** Representative time-lapse images showing a collapsing diplotene nucleus. Maximum-intensity projections were scaled identically. The frame showing initial contact between NE and SC was empirically identified. Note the asymmetric distribution of ZYG-12::GFP at the NE prior to and during collapse. Time stamps are min:sec. Scale bar, 5 µm. **g.** Another example of a late pachytene/early diplotene nucleus as it undergoes collapse, displayed as temporal projections of 3 successive timepoints to highlight the shrinking of NE without concomitant contraction of SC from 40sec to 50sec. Time stamps are min:sec. Scale bar, 2 µm. **h.** Quantification of NE and SC sizes measured from time-lapse images. 10 collapsing nuclei from 8 animals were measured. The point of contact between NE and SC was used to align data, and the final frame was used to for normalization. Mean ± SD are plotted. **i.** Quantification of the rate of nuclear collapse both before (‘pre-’) or after (‘post-’) contact between NE and SC. Medians (black crossbars) and means (black boxes) are shown. *p*-values calculated using Mann-Whitney test.

During late prophase in LMN-1-depleted germlines, we observed striking relocalization of SUN-1 and ZYG-12 to form an asymmetrical cap on the nuclei, shortly preceding nuclear collapse (**Figure 2f**, **3a; Supplementary Figure 8; Supplementary Video 4**). These NE also often showed obvious invaginations, indicative of active force being exerted at these sites. A reduction in nuclear size preceded the reduction in chromosomal volume, with the rate of NE collapse exceeding that of the chromosomal compaction before they make contact (**Figure 2f-i; Supplementary Videos 4,5**). This indicates that the collapse is driven by destabilization of the nuclear envelope and associated proteins, rather than by compaction of the chromosomes.

**Figure 3:**
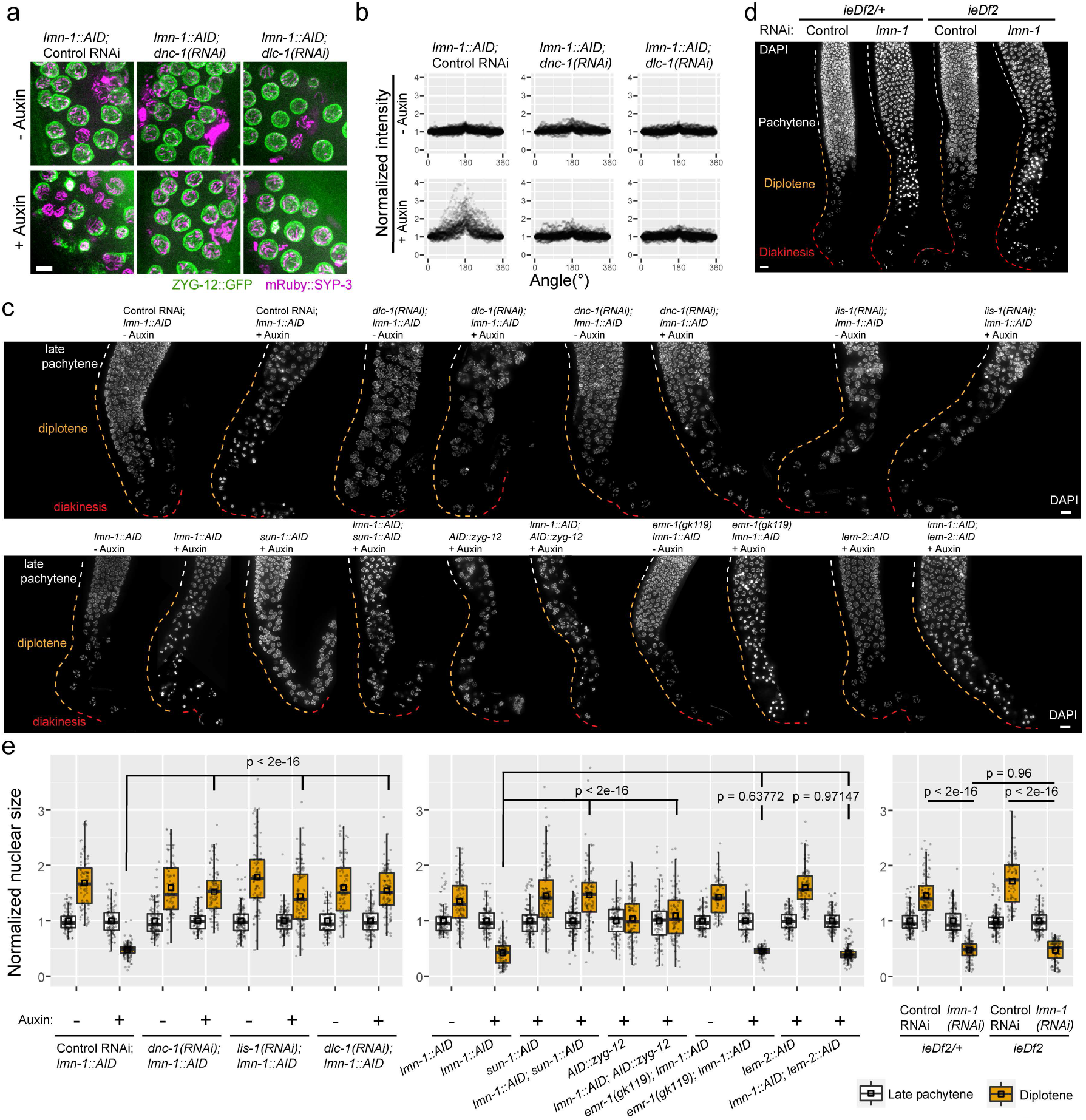
Nuclear collapse is rescued by disrupting dynein or LINC complex, but not the connections between LINC complex and pairing centers. **a.** Composite images showing mRuby::SYP-3 (magenta) and ZYG-12::GFP (green) at the NE of diplotene nuclei of indicated genotypes or treatments. All images are maximum-intensity projections and were scaled identically. Scale bar, 5 µm. **b.** Quantification of ZYG-12::GFP fluorescence intensities as a function of angle at the NE of diplotene nuclei of indicated genotypes or treatments. Sum of fluorescent intensity after background subtraction of ZYG-12::GFP from line-scan profiles along the circumference of each nucleus are mapped to a 0-360° angle range on a circle. The coordinate of maximum-intensity was aligned to 180° and the intensity at 0° was normalized to one (See **Supplementary Figure 7**). Unpaired two-sample two-sided *t*-test was used to calculate *p*-values comparing normalized peak intensities from 170 – 190° between each condition: p < 2.2e-16 between Control RNAi -/+ Auxin (66.42% increase in mean when +Auxin), p = 0.2603 between *dnc-1(RNAi)* -/+ Auxin (1.26% decrease in mean when +Auxin), p = 2.716e-06 between *dlc-1(RNAi)* -/+Auxin (3.85% increase in mean when +Auxin). Also see **Statistical Source Data**. **c.** Nuclear morphology at later stages of meiotic prophase. All worms were homozygous for *P_sun-1_::TIR1*. Colored dashed lines mark meiotic stages based on the anatomical positions of gonads from control animals. Scale bar, 10 µm. **d.** Nuclear morphology at later stages of meiotic prophase, from gonads dissected from animals of indicated genotypes or treatments. Pairing center proteins are present in *ieDf2/+* heterozygotes but absent in homozygotes. Colored dashed lines mark meiotic stages based on their distributions in controls. Scale bar, 10 µm. **e.** Normalized nuclear size from worms of indicated genotypes or treatments. Medians (black crossbars) and means (black boxes) are shown. One-way ANOVA and *post hoc* pairwise *t*-tests were used to compute the *p*-values (adjusted by the Benjamini-Hochberg method). The colors correspond to dashed lines in (**c**) and (**d**).

### Dynein-mediated forces promote LINC complex polarization and nuclear collapse

Movement of LINC complexes in early meiosis is driven at least in part by their interaction with dynein motors moving along microtubules in the cytoplasm (Wynne et al., 2012; Zhou et al., 2009). We found that depletion of either dynein light chain DLC-1 or the dynein activator dynactin (DNC-1) by RNAi largely eliminated the asymmetric localization of LINC factors during late prophase (**Figure 3a,b; Statistical Source Data**). Diplotene collapse was also rescued by co-depletion of DNC-1, DLC-1 or LIS-1 (another dynein activator), but not DYLT-1, which is dispensable for some dynein functions (Liu et al., 2015; O’Rourke et al., 2007) (**Figure 3c,e; Supplementary Figure 9**). Co-depletion of dynein heavy chain (DHC-1) or light intermediate chain (DLI-1) also prevented apparent collapse in diplotene, although nuclear positioning was markedly perturbed (**Supplementary Figure 9**). Co-depletion of SUN-1 or ZYG-12 using auxin-inducible degradation (**Supplementary Figure 1b,10,11**) also rescued nuclear collapse, further indicating that this process is driven by asymmetric forces acting on nuclei through the LINC complex (**Figure 3c,e**). Notably, co-depletion of SUN-1 did not suppress the increased and persistent RAD-51 foci in animals depleted of LMN-1 and SPO-11, reinforcing the conclusion that SPO-11 independent DNA damage does not trigger collapse (**Supplementary Figure 4**). We also introduced a deletion of the genes encoding ZIM-1, ZIM-2, ZIM-3, and HIM-8, four zinc finger proteins that bind to pairing centers and connect them to the LINC complex during early meiosis(Harper et al., 2011). This deletion did not rescue the nuclear collapse caused by LMN-1 depletion, indicating that asymmetric forces acting on late meiotic nuclei do not require attachment of LINC complexes to pairing centers (**Figure 3d,e**). However, this does not rule out a potential role for chromosome tethering to the NE through other sites or mechanisms (Gerstein et al., 2010; Ikegami et al., 2010).

### The lamin meshwork cooperates with the inner nuclear membrane proteins SAMP-1 and EMR-1/LEM-2 to stabilize oocyte nuclei

If mechanical force transmitted via the LINC complex drives nuclear collapse, conditions that disrupt LINC complex function should rescue this effect. We thus tested the effects of depleting other INM proteins implicated in LINC complex function. These included emerin (EMR-1), a homolog of which is required to recruit a KASH protein to the NE in *Drosophila* (Mandigo et al., 2019), LEM-2, since its fission yeast ortholog interacts with the SUN protein Sad1 (Hiraoka et al., 2011), and SAMP-1, whose mouse ortholog is a component of the transmembrane actin-associated nuclear (TAN) lines that contain the SUN2/nesprin-2G LINC complex (Borrego-Pinto et al., 2012; Gudise et al., 2011). SAMP-1 in *C. elegans* is also required for mechanotransduction during somatic nuclear migration (Bone et al., 2014). Surprisingly, meiosis and germline nuclear morphology appeared normal in the absence of any of these three proteins, and their depletion did not rescue nuclear collapse caused by LMN-1 depletion (**Figure 3c,e;Figure 4a,b; Supplementary Figure 11, 12**). Indeed, depletion of SAMP-1 exacerbated the effects of LMN-1 depletion, in that collapsed nuclei were observed in a more distal (earlier) region of the germline, as early as meiotic entry (**Figure 4a,b**). Because LEM-2 and EMR-1 have partially overlapping functions (Morales-Martínez et al., 2015), we tested the effects of co-depleting them, and found that this also sensitized nuclei to LMN-1 depletion (**Figure 4c,d**). LINC complexes also redistributed asymmetrically in nuclei undergoing precocious collapse upon co-depleting LMN-1 and EMR-1/LEM-2 (or SAMP-1) (**Supplementary Figure 13; Statistical Source Data**). Co-depletion of SUN-1 rescued the earlier nuclear collapse, indicating a common cause of meiotic nuclear collapse during oogenesis (**Figure 4**). Thus, we conclude that EMR-1, LEM-2 and SAMP-1 play LMN-1-independent roles in stabilizing nuclei against external forces, which become critical when LMN-1 is absent. Taken together, our findings illustrate that meiosis and oogenesis involve forces that can destabilize cell nuclei, and that the nuclear lamina and a network of inner nuclear envelope proteins both contribute to resisting these forces to enable successful reproduction (**Figure 5**).

**Figure 4:**
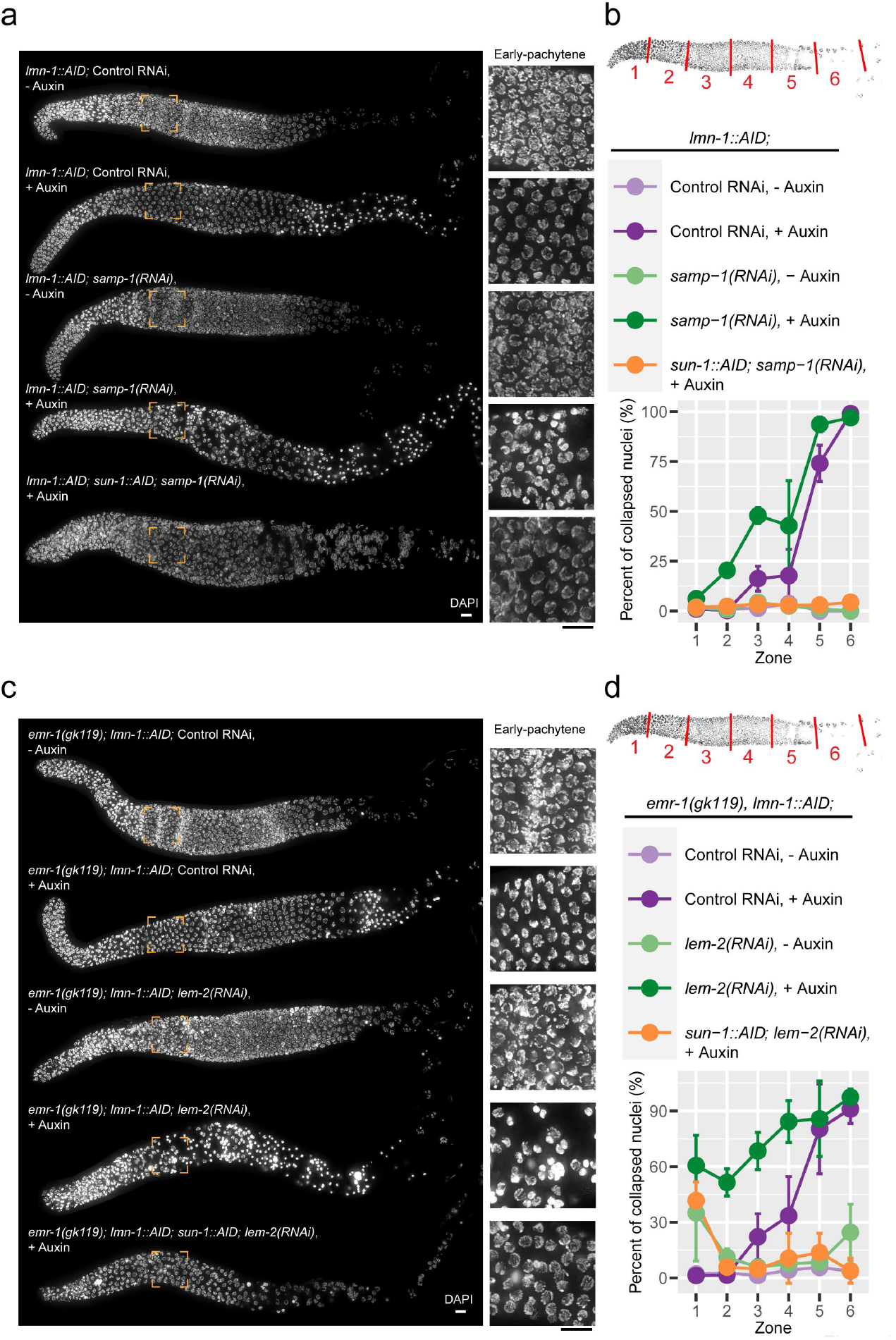
Nuclear collapse due to lack of LMN-1 is exacerbated by co-depletion of SAMP-1 or EMR-1/LEM-2. **a** and **c**. Nuclear morphology throughout meiosis. All worms were homozygous for *P_sun-1_::TIR1* or *P_gld-1_::TIR1* (omitted in the figure). Scale bars, 10 µm. Insets for early pachytene are 3× magnified. **b** and **d**. Fraction of collapsed nuclei as a function of meiotic progression. Schematics at the top shows distal gonad being divided into six zones of equal lengths, so that the percent of nuclei with collapsed/condensed chromatin in each zone can be quantified. Three worms were measured per condition. Mean ± SD are plotted. Pairwise comparisons for proportions were used to compute the *p*-values (adjusted by the Benjamini-Hochberg method, see **Statistical Source Data**).

**Figure 5:**
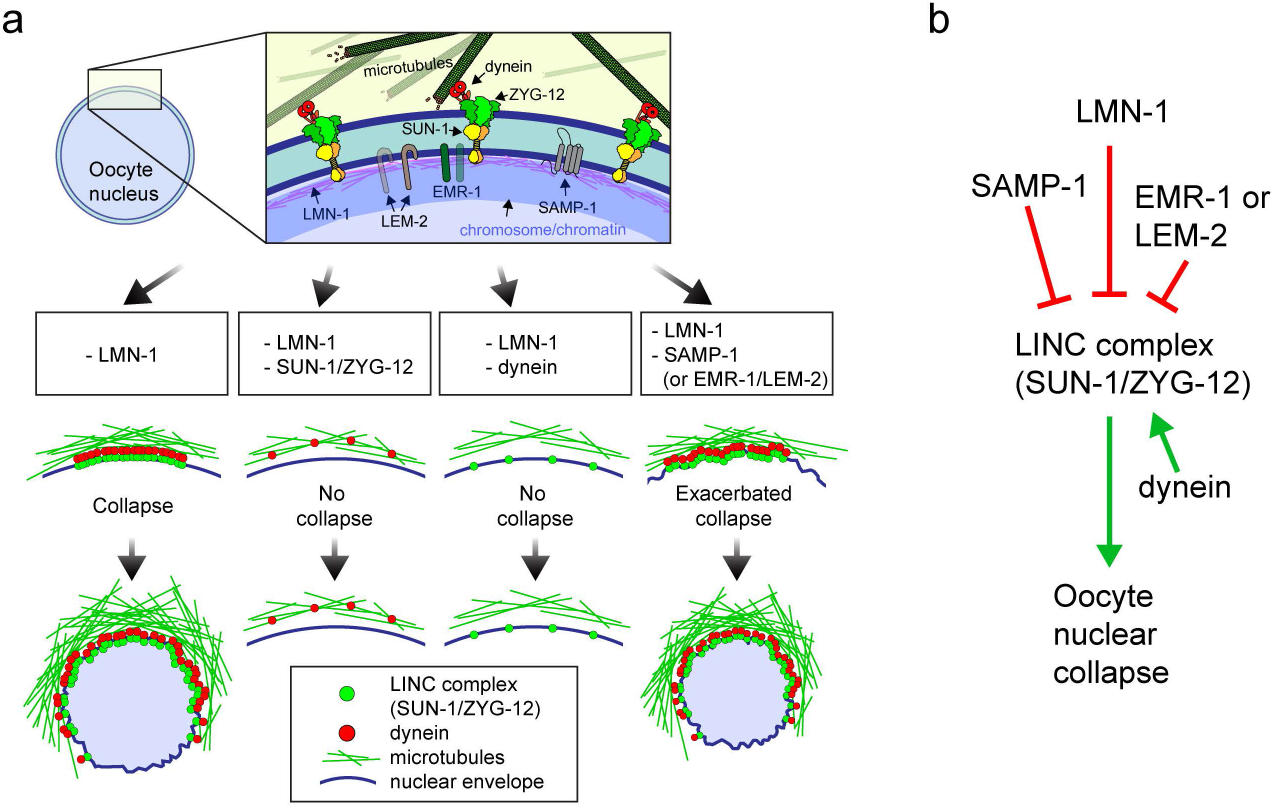
A cooperative network stabilizes oocyte nucleus against mechanical forces. **a.** Illustration of components influencing nuclear stability during oogenesis. LMN-1 (magenta) and INM proteins (EMR-1, LEM-2 and SAMP-1) likely provide mechanical rigidity at INM that withstand forces generated by MT/dynein and transmitted by the LINC complex to the NE. Depletion of LMN-1 leads to polarization of LINC complexes on one side of the nucleus, transmitting unbalanced force leading to nuclear collapse. Co-depletion of SUN-1 or ZYG-12, or components/regulators of the dynein motor rescues nuclear collapse; Simultaneous depletion of EMR-1, LEM-2 or SAMP-1 exacerbates nuclear collapse. **b.** Summary of forces contributing to nuclear stability during *C. elegans* oogenesis. Forces inhibiting collapse are depicted with red arrows, while those that promote collapse are depicted with green arrows.

## DISCUSSION

What determines the timing of nuclear collapse? LMN-1-deficient nuclei collapse as oocytes undergo massive growth and cellularization, which lead to hydrodynamic destabilization (Chartier et al., 2021; Geles and Adam, 2001). However, depletion of DNC-1 rescues nuclear collapse without impeding oocyte growth; similarly, depletion of OOC-5 rescues collapse without obvious effects on oocyte nuclear volume (Huelgas-Morales et al., 2020), so growth alone is not sufficient to induce collapse; forces acting directly on the nuclear envelope are clearly important. In the absence of LMN-1 and SAMP-1 or EMR-1/LEM-2, nuclei collapse during early meiosis, likely due to a reduction in lamina stability at this stage: LMN-1 is phosphorylated during early meiosis and can be extracted from the NE by detergent exposure prior to fixation, likely reflecting reduced crosslinking (Link et al., 2018). In other organisms, lamins are replaced by more labile isoforms or completely dismantled during early meiotic prophase to enable chromosome movements that promote homolog pairing and synapsis (Dernburg, 2013). Thus, lamin-independent mechanisms likely play a conserved role in reinforcing the NE during meiosis.

Our finding that LMN-1 opposes mechanical forces mediated through LINC complexes seems at odds with previous observations that in migrating mouse fibroblasts, the lamina works together with the LINC complex during nuclear movement. In *LMNA* mutant cells, TAN lines containing LINC complex cannot be anchored properly in the NE and consequently “slip over” the nucleus instead of moving with it(Folker et al., 2011). However, it is consistent with our data that LMN-1 is required to prevent “unhinged” movement of LINC complex across the NE driven by the cytoskeleton. Our observations are also consistent with previous findings that a disease-causing *lmn-1* allele leads to dynein-dependent NE rupture during pronuclear migration in *C. elegans* (Penfield et al., 2018), that cell polarity defects in Hutchinson–Gilford progeria syndrome (HGPS) fibroblasts can be rescued by inhibiting MTs/dynein (Chang et al., 2019), and that NE rupture, viability and function of *Lmna* KO myotubes in mice can be rescued by reducing MT-based force via depleting a kinesin or expressing a dominant negative KASH (Earle et al., 2020). All of these observations support the idea that the nuclear lamina is important to stabilize nuclei by counteracting MT/LINC mediated forces.

Oocyte nuclei experience a range of unusual forces, including those arising from chromosome movements during meiosis, large-scale cell and nuclear growth during maturation, and extracellular forces both within the ovarian cortex and during ovulation. In mice, compressive forces from the surrounding granulosa cells are important to maintain the dormant state of oocytes arrested at the diplotene (dictyate) stage, through dynein based nuclear rotation (Nagamatsu et al., 2019). Given that LINC complexes are required for nuclear rotation(Lee et al., 2015; Wu et al., 2018), and our finding that lamina is required for anchoring LINC in place against dynein-based motility, it is plausible that nuclear lamina is important for maintaining the dormant state of primordial oocytes in mammals. Intriguingly, polymorphisms in the emerin-associated NE protein Nemp1, which confers stiffness to the NE, have been associated with early menopause in humans. Nemp1 is also essential for fertility across metazoans, including in *C. elegans* (Tsatskis et al., 2020), adding another component to the network of factors that contribute to nuclear envelope integrity in oocytes. Our work adds to a growing body of evidence that multiple pathways contribute to the resilience of oocyte nuclei during metazoan reproduction.

## METHODS

### Generation of worm strains

All *C. elegans* strains were maintained at 20°C under standard conditions (Brenner, 1974). New alleles generated in this study and their detailed information are listed in **Supplementary Table 2**. A complete list of *C. elegans* strains generated in this study is also provided in **Supplementary Table 3**. All new alleles in this study were generated using CRISPR/Cas9 genome editing (Alt-R CRISPR-Cas9 gRNA products, IDT), as previously described (Zhang et al., 2018). Briefly, duplexed gRNA-Cas9 RNP complex was injected into the gonad of young adults, with *dpy-10* co-CRISPR to facilitate the screen of edited progeny (Arribere et al., 2014; Paix et al., 2015). Custom designed gRNA against gene of interest (200µM, IDT; sequences can be found in **Supplementary Table 4**) was mixed with gRNA targeting *dpy-10* (50µM) at 3:1 (v:v) ratio, and then combined with tracrRNA (200µM, IDT) at 1:1 (v:v) ratio, annealed at 95°C for 5min followed by room temperature for 5min. The RNAs were then complexed with Cas9-NLS (40µM, UC Berkeley MacroLab) at room temperature for 5min. Repair templates, including dsDNA repair template (purified PCR product of gBlock from IDT) and ssDNA repair template for *dpy-10* (IDT) were added to the injection mix, which was centrifuged at 13200 rpm in 4°C for 10min prior to loading quartz needles (Sutter Instrument) for micro-injection (Eppendorf). 3∼4 injected hermaphrodites were pooled per plate and incubated at 20°C for 4 days, before individual F1 Dumpy or Roller worms were transferred to new plates. After 3 days at 20C, individual F1 hermaphrodite from each plate was lysed and screened with PCR genotyping. Both N- and C-terminal tagging of endogenous LMN-1 with the AID degron resulted in a loss-of-function phenotype. N-terminal tagging of SUN-1 with AID also resulted in 100% inviable progeny. C-terminal tagging of SUN-1 with AID did not disrupt SUN-1 function but the protein was refractory to auxin-inducible degradation. Simultaneous AID-tagging of LMN-1 and LEM-2 in *emr-1(gk119)* mutants resulted in embryonic lethality.

### Fertility and viability

Brood size, viability and male index among self-progeny were quantified as described (Zhang et al., 2018). Briefly, fresh OP50 plates were made with thin lawns, and L4 hermaphrodites (P0) were plated individually. Animals were transferred to a new plate every 24 hours for a duration of 4-5 days until egglaying ceased. Eggs were counted after removal of the parents, and viable progeny were counted 2-3 days later. To quantify brood size, viability and male self-progeny in the presence of auxin, age-matched hermaphrodites at L1 or L4 stages were transferred onto auxin plates following the same procedures. When counting eggs on non-auxin plates, only those eggs that appeared firm with intact eggshells were counted; when counting eggs on auxin plates, however, due to the many dead and deformed eggs, only eggs that have intact eggshells were counted.

### Auxin-induced degradation

Unless otherwise specified, hermaphrodites were picked at the L4 stage and transferred to plates containing 2mM indole acetic acid (IAA, auxin) starting at 12 hours after L4. Worms were left on auxin plates for 12 hours before dissection (for immunofluorescence) or live imaging (of acridine orange stained nuclei in live animals, or in vivo imaging of fluorescent proteins). Auxin plates were prepared as previously described (Zhang et al., 2018). An overnight culture of OP50 in LB was concentrated by 10-fold and supplemented with auxin before seeding the auxin plates. All experiments were performed at 20°C.

### RNA interference

RNAi interference experiments were carried out according to established protocols with minor modifications (Davies et al., 2018; Timmons et al., 2001). Briefly, individual HT115 bacterial clones were streaked from frozen stocks of *C. elegans* RNAi feeding library constructed by Julie Ahringer’s group (Kamath and Ahringer, 2003; Kamath et al., 2003) (see **Supplementary Table 5**) onto 10cm Luria broth (LB) agar plate supplemented with carbenicillin (final 100µg/mL) and tetracycline hydrochloride (topically spread, 2µL of 25mg/mL stock aqueous solution diluted in 400uL LB). A single colony was inoculated into 5mL LB supplemented with carbenicillin (final 50µg/mL) and tetracycline hydrochloride (final 12.5µg/mL) and cultured for about 16 hr overnight at 37°C, 250rpm. NGM agar plates for RNAi were made with the following supplements freshly added before pouring: carbenicillin (final 50µg/mL), tetracycline hydrochloride (final 12.5µg/mL), IPTG (final 1mM). Auxin was only added in plates for simultaneous RNAi and auxin-inducible degradation. These RNAi plates were seeded with 300µL of non-concentrated bacterial culture, allowed to dry at room temperature, and moved to a dark 30°C incubator for 48 hours before transferring worms onto them. All RNAi plates are covered in aluminum foil and kept at 4C once made for a couple of months (unseeded) or a couple of weeks (seeded). HT115 clones carrying L4440 empty vectors were used as RNAi controls. For RNAi experiments, L4 hermaphrodites were transferred onto RNAi plates and dissected 48 hours later; when combined with inducible degradation, animals were transferred to plates containing both IPTG and auxin for 12hr before dissection. All experiments were performed at 20°C.

### Immunofluorescence

Immunofluorescence was carried out as previously described (Phillips et al., 2009) with minor modifications. Dissected gonads were transferred to low-retention microcentrifuge tubes, and subsequent steps (fixation, permeabilization, blocking and staining etc.) were performed in suspension. The following antibodies were used: anti-V5 (mouse, P/N 46-0705, Invitrogen), anti-HA (mouse monoclonal, #26183, Invitrogen), anti-GFP (mouse monoclonal, Roche, 1:400), anti-SYP-1 (goat, affinity purified, 1:400 (Harper et al., 2011)), anti-HTP-3 (chicken, 1:400 (MacQueen et al., 2005)), anti-HIM-8 (rat, 1:400 (Phillips et al., 2005)), anti-NPP-7 (rabbit polyclonal, SDIX, SDQ0870, 1:1,000), anti-NPP-10 (rabbit polyclonal, SDIX, SDQ0828, 1:1,000), anti-RAD-51 (rabbit polyclonal, SDIX, Cat#29480002, 1:20,000 (Zhang et al., 2018)) and secondary antibodies conjugated to Alexa 488, Cy3, or Cy5/AF647 (Jackson ImmunoResearch or Life Technologies; 1:400).

All images of fixed samples were acquired using a DeltaVision Elite system (GE) equipped with a 100X 1.4 NA oil-immersion objective (Olympus). Wide-field images were deconvolved using the SoftWoRx Suite (Applied Precision, GE). To compare between individual gonads or slides, the same exposure times were used for all samples. All fixed gonads were imaged as 3D image stacks at intervals of 0.2 µm. Maximum-intensity projections of deconvolved 3D images were assembled using Fiji and Illustrator (Adobe) for figures, unless otherwise noted.

### Acridine orange assay

Acridine orange staining and washing were carried out according to established protocols with minor modifications(Gartner et al., 2004). The assay was timed so that the entire duration of auxin treatment was consistent with the other AID experiments (*i.e*., the time required for acridine orange staining and washing was included in the total time of auxin treatment). Briefly, acridine orange staining solution was prepared by diluting a 10 mg/ml stock solution (Molecular Probes A-3568) 1:200 into M9 buffer. Age-matched adult hermaphrodites from control or auxin plates were picked to the middle of OP50 lawn of fresh control and auxin plates. 900uL staining solution was gently pipetted onto each plate until the entire surface (including the OP50 lawn) was covered. Animals remained on top of the OP50 lawn to ensure they have active feeding behavior, which is important for the intake of acridine orange. Plates were then covered with aluminum foil to shield them from light for one hour at room temperature without disturbance. Worms were then washed off the plate with 1.5mL M9 buffer and transfer to low-retention 1.5mL tubes (Fisherbrand) and centrifuged for 10 sec in a tabletop mini centrifuge. The supernatant was removed and replaced with M9 buffer. Centrifugation and washing were repeated twice more. After the final wash, the worms were transferred to new control and auxin plates and kept in the dark for 45 minutes at room temperature before mounting on agarose pads. They were imaged using a spinning disc confocal microscope equipped with differential interference contrast (DIC) and 488nm/561nm excitation laser to image and score acridine orange positive meiotic nuclei (indicative of apoptosis) within one hour (see ‘Live imaging’). To prevent desiccation of animals during live imaging, about four worms were picked onto an agarose pad at a time. Low retention pipette tips (Fisherbrand) were used throughout the washing process.

### Live imaging

For live imaging, gravid hermaphrodite adults were picked into a 10-µL drop of water containing 250 µM tetramisole hydrochloride on a freshly prepared agarose pad (7.5% in water). The immobilized worms were overlaid with #1.5 coverslips (0.16 to 0.19mm, #152222, SLIP-RITE, Richard-Allan Scientific), sealed to the slide using VALAP (1:1:1 [w:w:w] vaseline:lanolin:paraffin) and imaged immediately. Image acquisition was carried out on a Marianas spinning-disc confocal microscope (Intelligent Imaging Innovations, Inc.) at ambient temperature (21°C), using a 100X 1.46 NA oil immersion objective. To analyze LINC complex mobility, 3D image stacks at specific gonad zones/meiotic stages were acquired every 5 sec for a duration of 30 to 60 time points, with 10 Z-sections at intervals of 0.5 µm per time point. To examine the dynamics of diplotene nuclear collapse, image stacks were acquired every 10 s. Differential interference contrast (DIC) was used to facilitate identifying gonad zones, and 488 or 561 excitation lasers were used at identical parameters per set of experiment for both control and experimental groups. Live imaging following previously described protocols using 100 nm polystyrene beads (Polysciences, cat#00876) and serotonin creatinine sulfate (Sigma) was also carried out with no observable differences for the short-term imaging used in this work(Rog et al., 2017).

### Image analysis

To quantify nuclear volume (**Supplementary Figure 3**) measured with the DAPI signal, the “Surface” function in Imaris (x64, 9.2.0, Bitplane) was used for 3D segmentation and quantification. All 3D segmentation was carried out while crossreferencing to NE staining to confirm segmented DAPI signals belong to the same nucleus (**Supplementary Figure 2,3**).

Quantification of the distribution of LINC complexes at the NE was performed using ‘freehand line’ and ‘plot profile’ tools in Fiji. Motion of of LINC complex patch/foci in 3D was analyzed in Imaris (x64, 9.2.0, Bitplane) using the “Spots” function for tracking and reference frames for drift correction. The quantification of shrinking dynamics of collapsing NE and SC was carried out in Fiji and shrinking speeds were computed using linear model in RStudio.

For quantifying the percentage of pairing and complete synapsis as a function of meiotic progression, a distal gonad was divided into five equal-distanced zones from premeiotic (distal) end to early diplotene (in **Supplementary Figure 4c,6c**). X-chromosome pairing was scored based on the number of HIM-8 foci per nucleus, and complete synapsis was scored by colocalization of SYP-1 and HTP-3. For quantifying the percentage of collapsed nuclei as a function of meiotic progression, a distal gonad was divided into six equal-distanced zones from the distal tip to early diakinesis (**Figure 4b,d**). Collapsed nuclei were scored based on smaller sizes and hypercondensed chromatin compared to adjacent nuclei in the same zone from the same animal as well as nuclei in the same zone from control animals without RNAi or auxin treatment.

### Statistics

All statistical analyses were carried out in RStudio (Version 1.2.5033), as noted in individual figure legends. All two-sample tests are unpaired and two-sided. To quantify asymmetry of LINC complex at diplotene, mean normalized peak intensities about 180° (170-190°) were compared using the two-tailed *t*-test (**Figure 3b**). To quantify LINC complex asymmetry at early pachytene NE (**Supplementary Figure 13b,d**), normalized intensity across 0-360° were combined as 0-180° due to the symmetric distribution about 180°, and the data were fit to a linear model (2-way ANOVA). Additional output of statistical tests performed in this study can be found in **Statistical Source Data**.

## Supporting information

Supplementary Statistics Data

## DATA AVAILABILITY

Source data are provided with this paper. All other data supporting the findings of this study are available from the corresponding authors upon reasonable request.

## ACKNOWLEDGEMENTS

We thank members of the Dernburg Laboratory for insightful discussions and critical reading of the manuscript. We thank Tim Davies (Durham University), Sophia Hirsch and Julie Canman (Columbia University) for advice on RNAi. We also thank Gregg Gundersen (Columbia University) and Ofer Rog (The University of Utah) for discussions, and Yu He (Yale University) for insights on quantitation. We are grateful Gabriela Huelgas-Morales and David Greenstein (University of Minnesota) for sharing related findings prior to publication. Some *C. elegans* strains used in this work were provided by the Caenorhabditis Genetics Center funded by the NIH - Office of Research Infrastructure Programs (P40 OD010440). This work was supported by a fellowship from the Life Sciences Research Foundation to CL (Grant Number 45283), a SURF fellowship to ZL, and support to AFD from the Howard Hughes Medical Institute.

## AUTHOR CONTRIBUTION

Chenshu Liu: Conceptualization, Funding acquisition, Data curation, Formal analysis, Investigation, Writing—original draft, Writing—review and editing

Zoe Lung: Funding acquisition, Data curation, Investigation; John S. Wang: Data curation, Investigation;

Fan Wu: Data curation, Investigation;

Abby F. Dernburg: Conceptualization, Funding acquisition, Investigation, Project administration, Writing—review and editing.

## DECLARATION OF INTERESTS

The authors declare no competing interests.

## SUPPLEMENTARY FIGURE LEGENDS

**Supplementary Figure 1: (Related to Figure 1 and 3).**
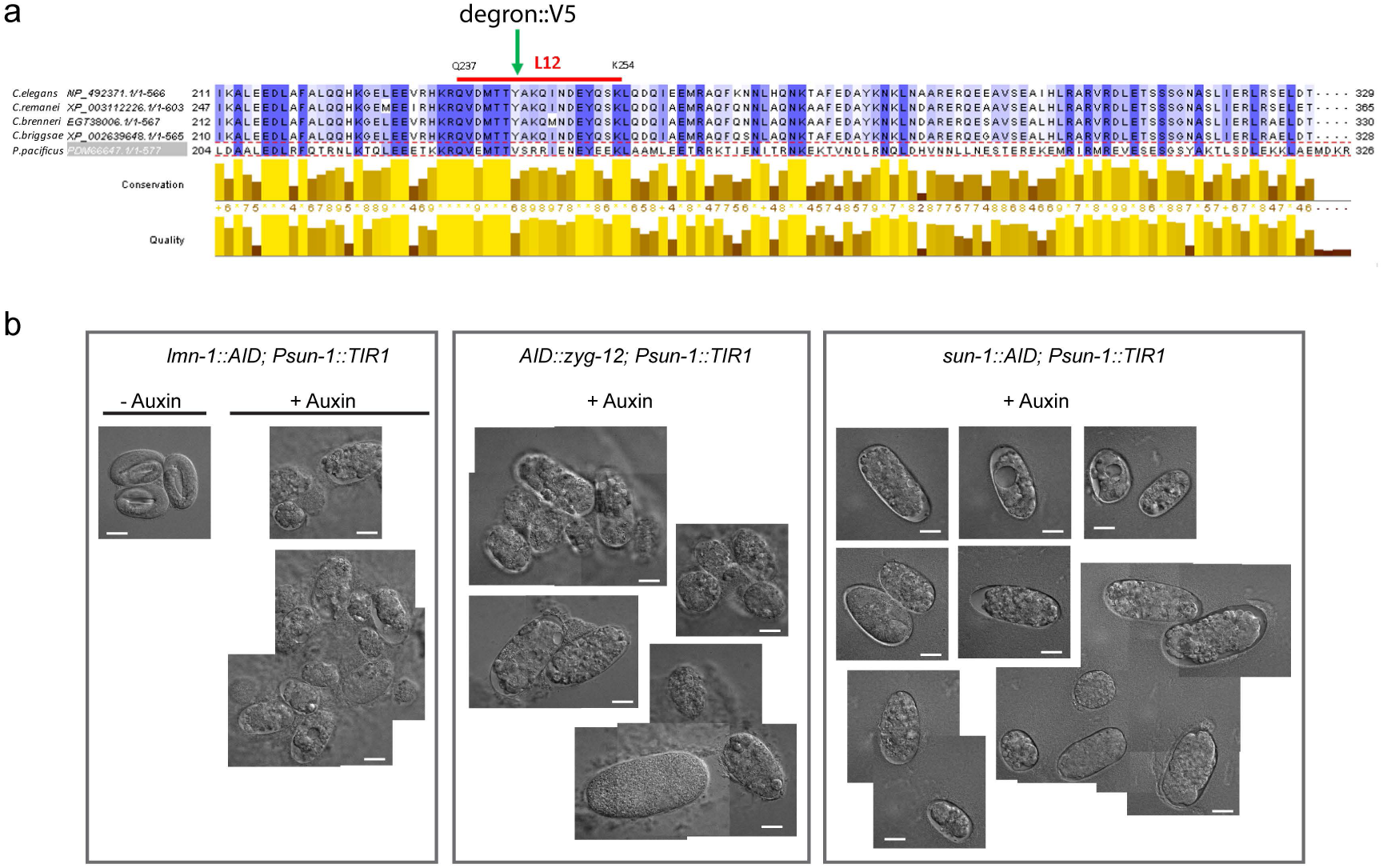
CRISPR tagging and auxin-inducible degradation of LMN-1 in *C. elegans*. **a.** Multiple sequence alignments of nematode LMN-1 proteins were generated using Clustal Omega and visualized using Jalview, showing position of degron/V5 insertion in *C. elegans* LMN-1. **b.** Morphology of eggs/embryos laid by hermaphrodite worms treated ± auxin for 48 hrs (since young adulthood). Images acquired using DIC. Scale bar, 20 µm.

**Supplementary Figure 2: (Related to Figure 1).**
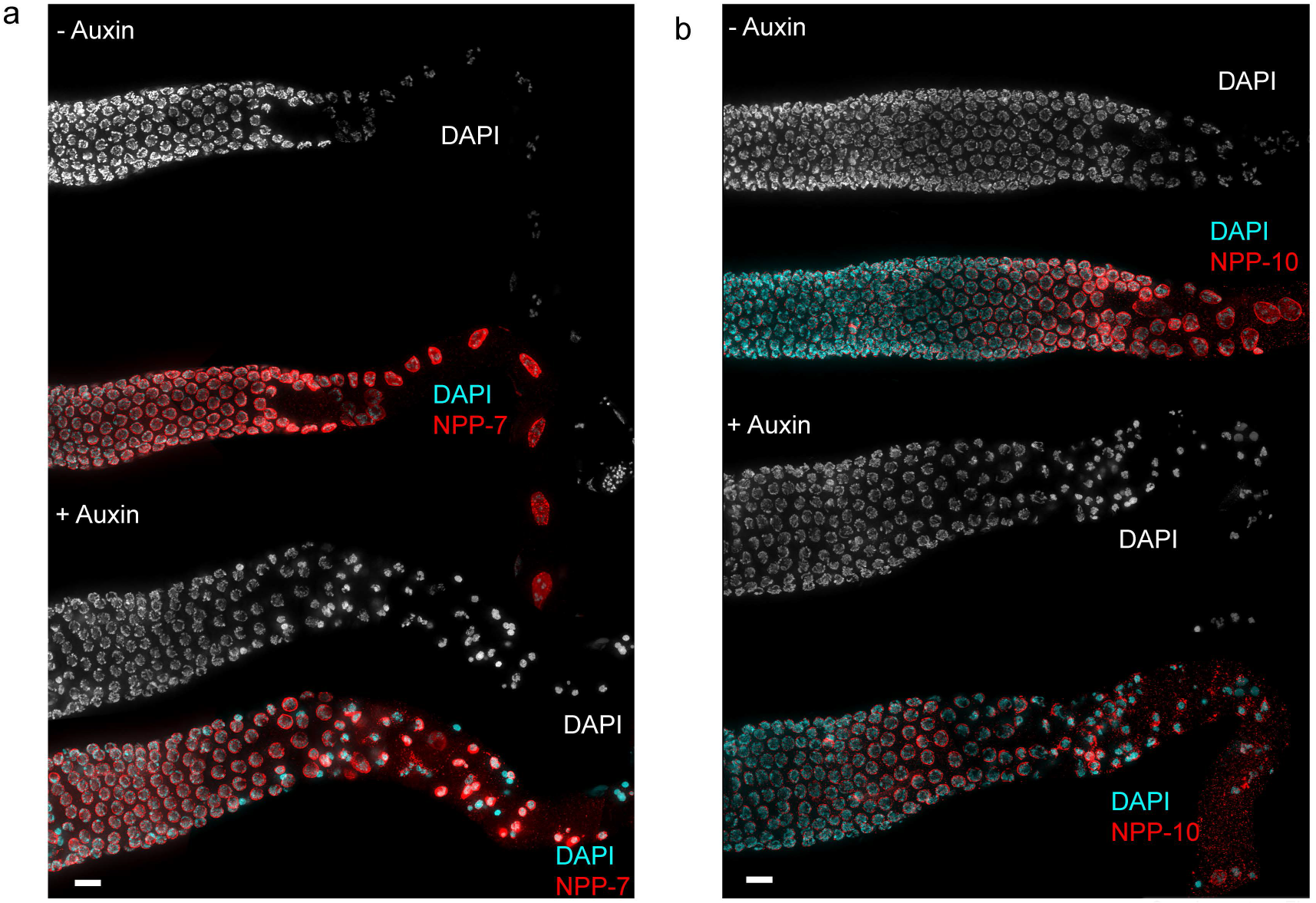
Representative images showing co-staining of DAPI and nucleoporins NPP-7 (a) or NPP-10 (b). Composite images are maximum-intensity projections. Meiosis progresses from left to right. Scale bars, 10 µm.

**Supplementary Figure 3: (Related to Figure 1).**
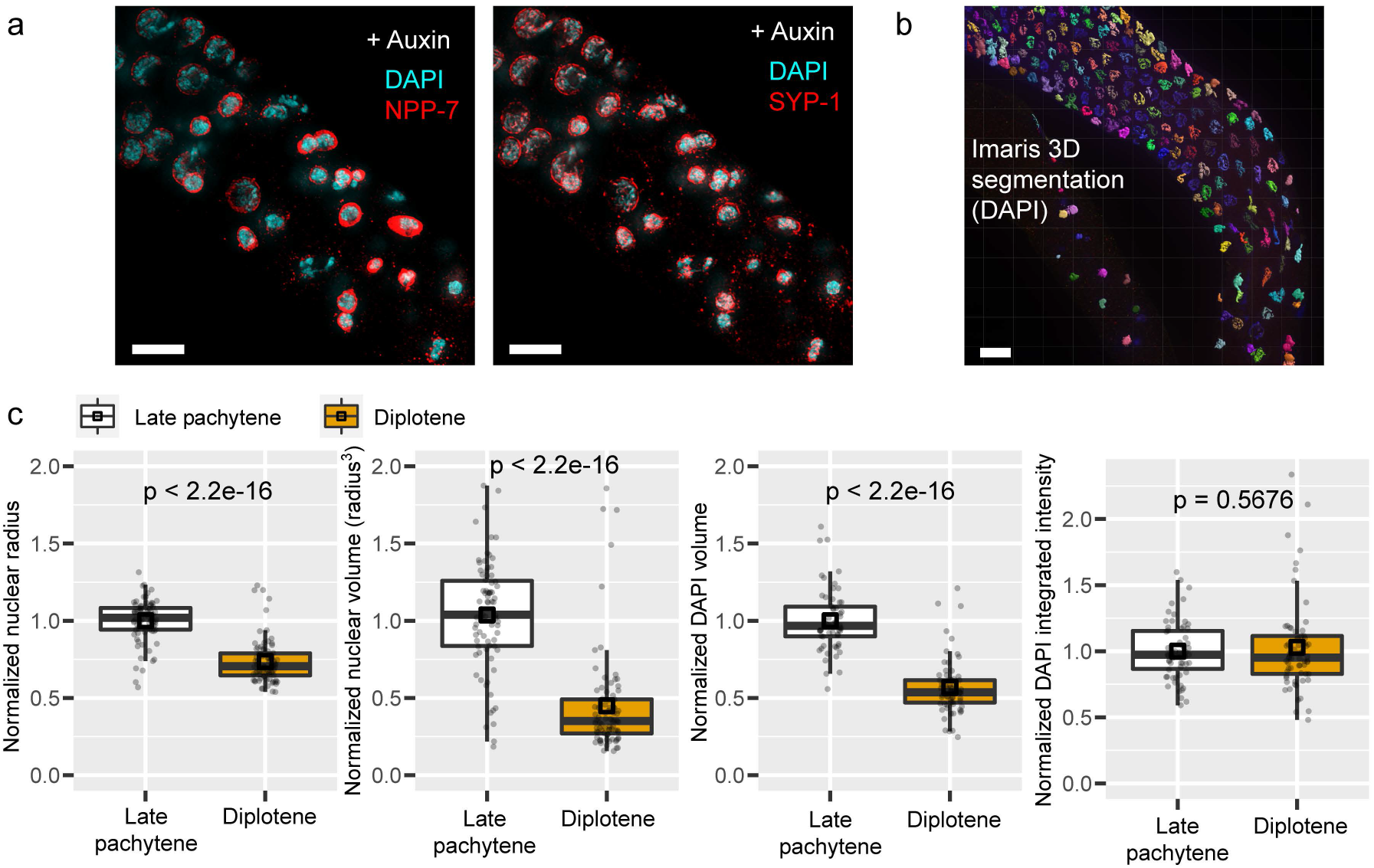
Quantifying nuclear collapse. **a.** Composite images (maximum-intensity projections) showing nuclei in late meiotic prophase stained for DAPI (cyan), NPP-7 (red) or SYP-1 (red). Meiosis progresses from top left to bottom right. Scale bars, 10 µm. **b.** Segmentation of chromosome volume based on DAPI fluorescence. Meiosis progresses from top left to bottom right. Scale bar, 10 µm. **c.** Quantification of nuclear radii based on NPP-7 or SYP-1 staining (with maximum Z projection), nuclear volume based on radius^3^, nuclear size based on DAPI volume and integrated DAPI fluorescence intensity per nucleus. Nuclear sizes in late pachytene were normalized to one. Unpaired two-sample two-sided *t*-test was used to calculate *p*-values. At least 58 nuclei from three animals were analyzed per stage.

**Supplementary Figure 4: (related to Figure 1 and 3).**
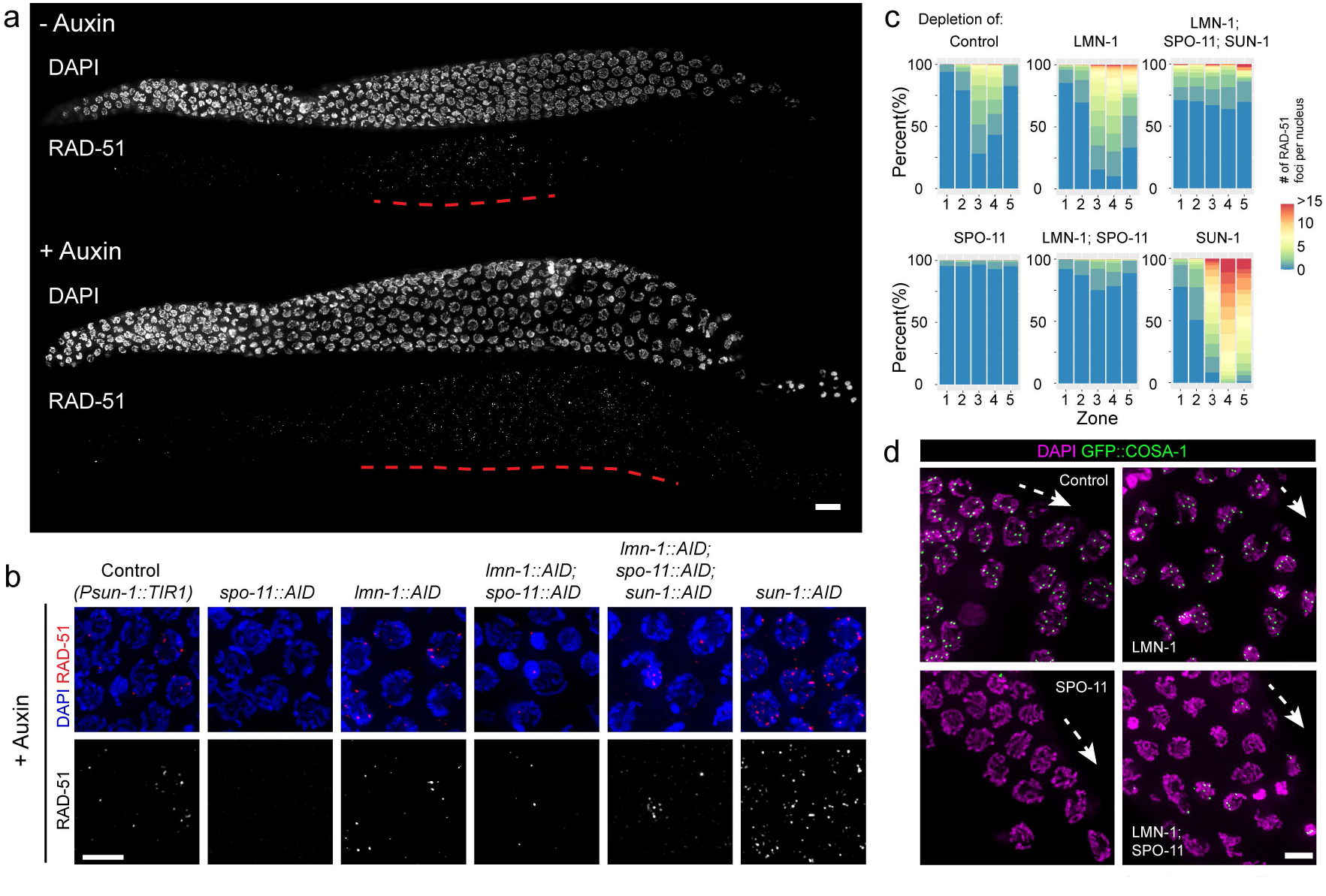
LMN-1 depletion causes SPO-11 independent DNA damage. **a.** LMN-1 depletion causes persistent DNA damage marked by RAD-51 foci. Scale bar, 10 µm. **b.** RAD-51 staining in mid-pachytene nuclei from control and hermaphrodites depleted of SPO-11, LMN-1, LMN-1 and SPO-11 both, or LMN-1 and SPO-11 and SUN-1 simultaneously. Control hermaphrodite has TIR1 but no AID-tagged genes. Animals were exposed to auxin for 24 hours from the L4 stage to young adulthood prior to dissection. Scale bar, 5 µm. **c.** Quantification of RAD-51 foci per nucleus as a function of meiotic progression. Gonads were divided into five zones of equal length spanning the premeiotic region to early diplotene (as in **Supplementary Figure 5**). Control, N = 964 nuclei (4 animals); SPO-11 depletion, N = 1480 nuclei (4 animals); LMN-1 depletion, N = 1781 nuclei (6 animals); LMN-1, SPO-11 double depletion, N = 1329 nuclei (5 animals); LMN-1, SPO-11, SUN-1 triple depletion, N = 844 nuclei (5 animals); SUN-1 depletion, N = 1201 nuclei (4 animals). Pairwise comparisons for proportions were performed to compute the *p*-values (adjusted by the Benjamini-Hochberg method, see **Statistical Source Data**). **d.** Designated crossover sites marked by GFP::COSA-1 foci in late prophase nuclei. Following LMN-1 depletion, nuclei still show six designated crossover sites, whereas GFP::COSA-1 foci were absent from apoptotic nuclei. Animals were exposed to auxin for 24 hours from the L4 stage to young adulthood prior to dissection. Dashed arrows indicate meiotic progression. Scale bar, 5 µm.

**Supplementary Figure 5 (Related to Figure 1 and 3).**
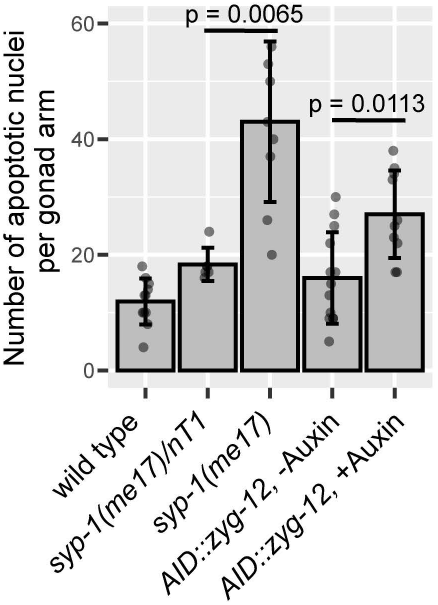
Germline apoptosis increases upon ZYG-12 depletion. Apoptosis was quantified using GFP::CED-1. *syp-1(me17)/nT1* heterozygotes and *syp-1(me17)* homozygotes were used as controls. Mean ± SD are plotted. *p*-values were calculated using the Mann-Whitney test.

**Supplementary Figure 6 (Related to Figure 2).**
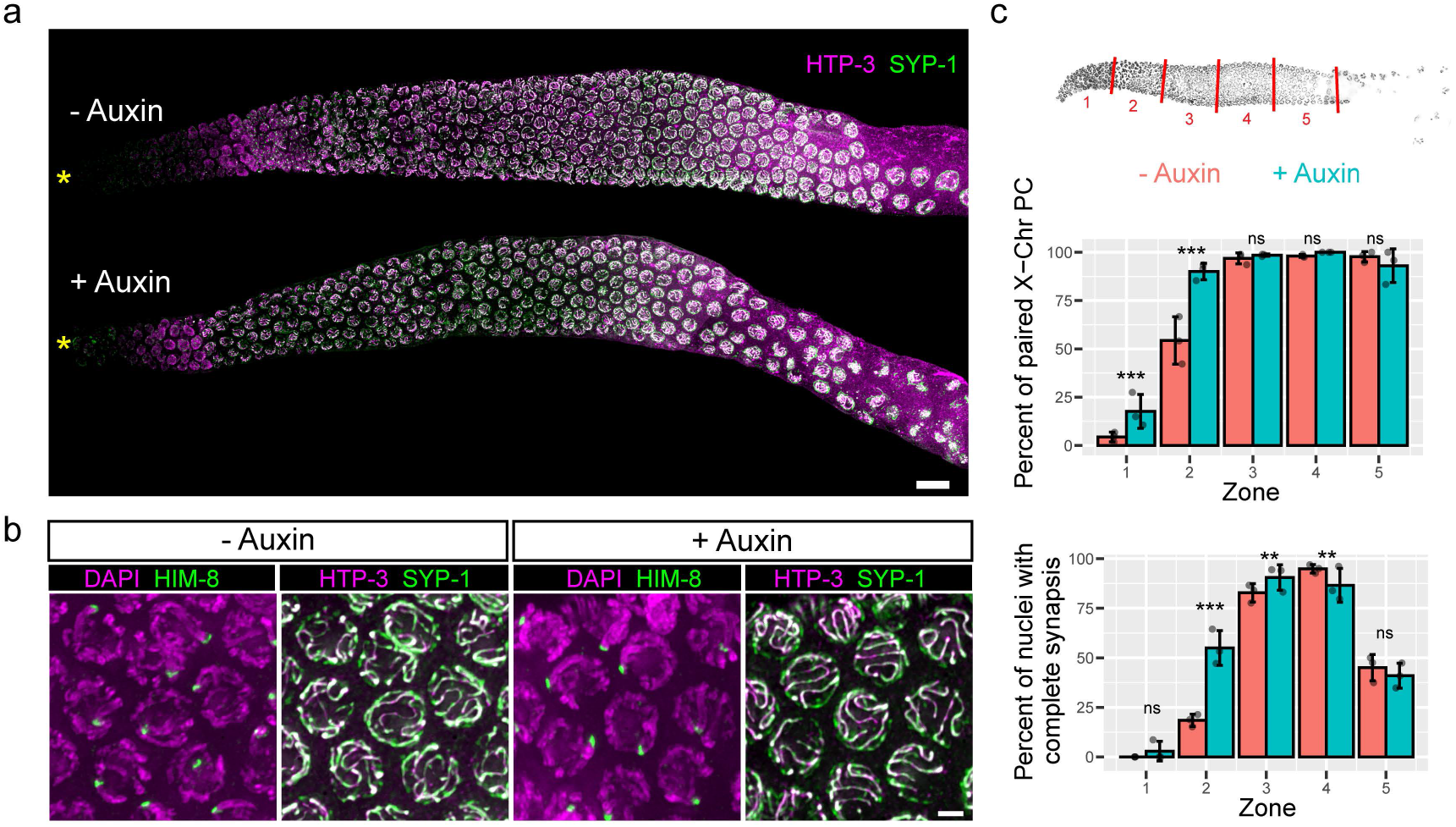
Chromosome pairing and synapsis occur normally following LMN-1 depletion. **a.** Dynamics of synapsis, based on immunostaining of SYP-1 and HTP-3. Yellow asterisks indicate the distal end of gonads. Scale bar, 10 µm. **b.** Normal pairing of X chromosomes is revealed by immunofluorescence of HIM-8, which localizes to the X-chromosome pairing centers. Scale bar, 2 µm. **c.** Quantification of homolog pairing and synapsis. Diagram of distal gonad divided into five zones of equal length. Three gonads were measured per condition. Mean ± SD are plotted. Two-sided two-proportions z-test was used for computing the p-values. ***, p ≤ 0.001; **, p ≤ 0.01; ns, p > 0.05 (see **Statistical Source Data**).

**Supplementary Figure 7 (Related to Figure 2).**
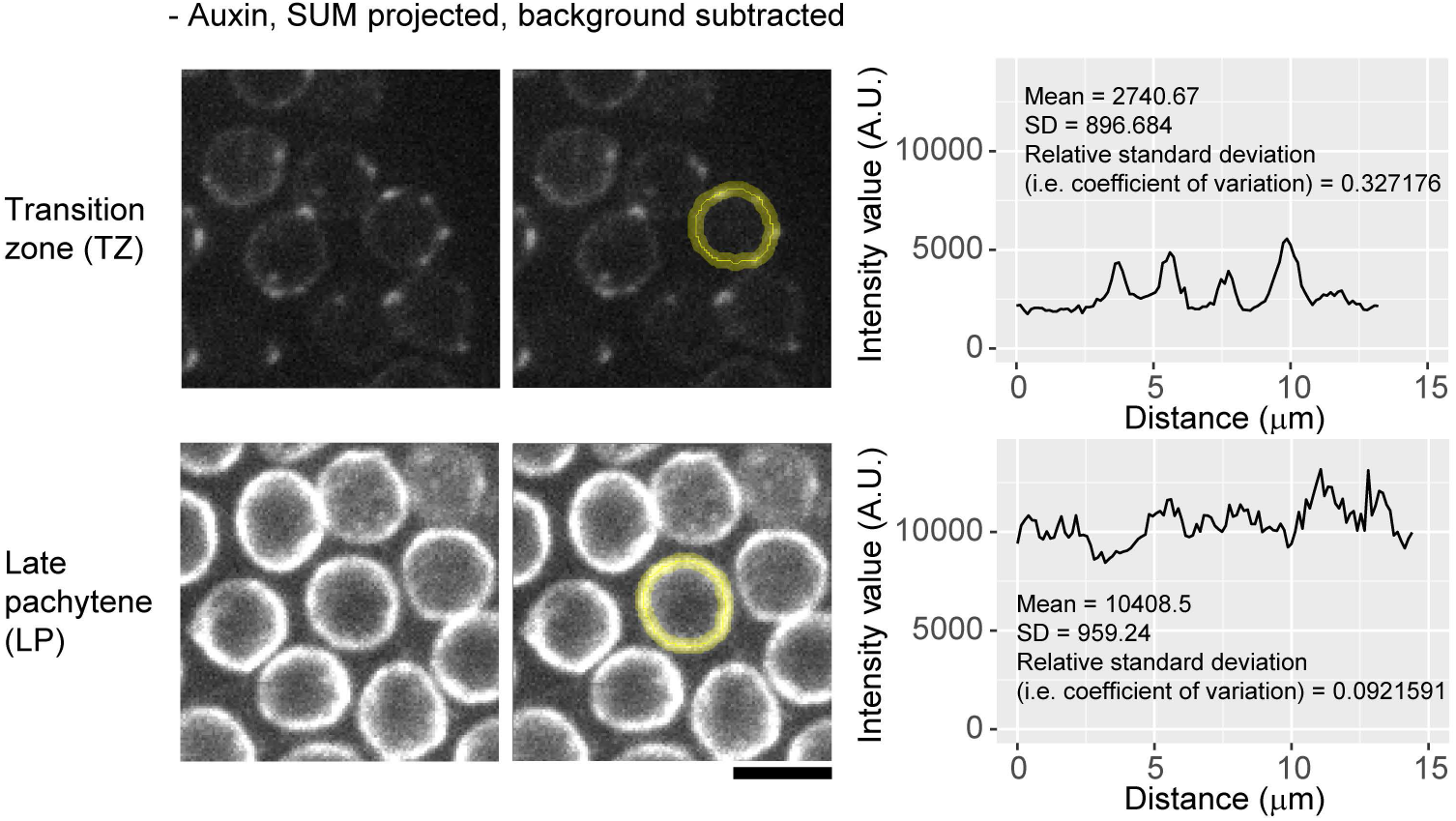
Quantifying SUN-1 distribution at the NE. Representative line-scan profiles of SUN-1::mRuby fluorescence intensity at the circumference of individual nuclei. Grayscale images are additive projections and are scaled using the same look up table (LUT). Scale bar, 5 µm.

**Supplementary Figure 8 (Related to Figure 3).**
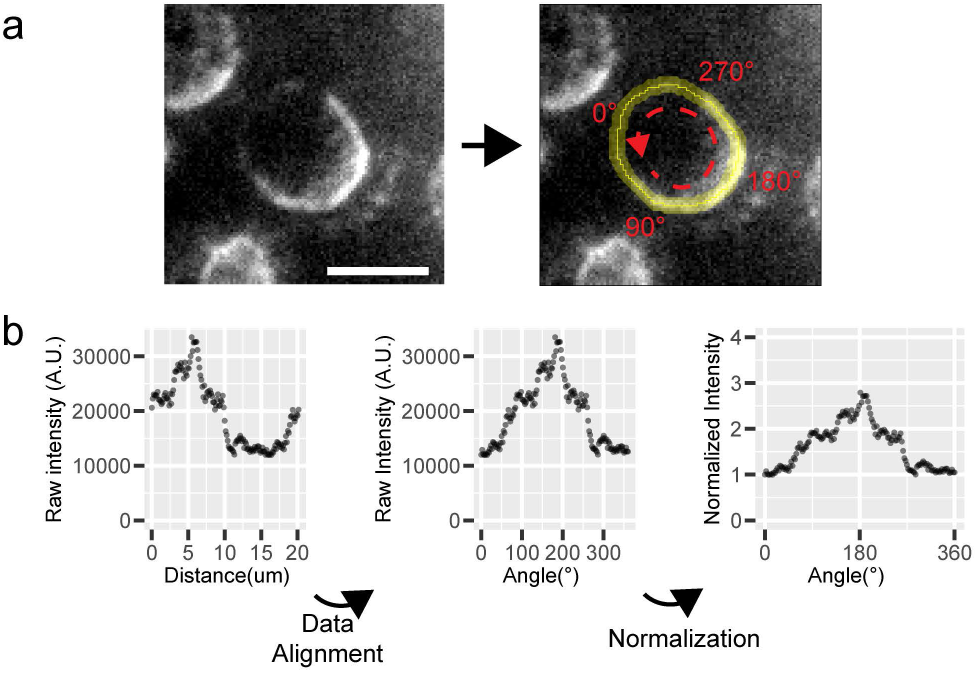
Quantifying asymmetric ZYG-12 distribution at the NE. **a.** Grayscale images of additive projections with background subtraction showing polarized ZYG-12::GFP distribution at the NE in a LMN-1 depleted diplotene nucleus. The right panel shows line scan profile along the circumference of the nucleus and approximate locations of angles mapped subsequently during data alignment. Scale bar, 5 µm. **b.** Data alignment and normalization. Raw intensity measurement from line-scan in (**a**) was aligned and mapped such that 180° corresponds to the coordinate along the NE’s circumference with the maximum-intensity. The intensity at 0° was normalized as one.

**Supplementary Figure 9 (Related to Figure 3).**
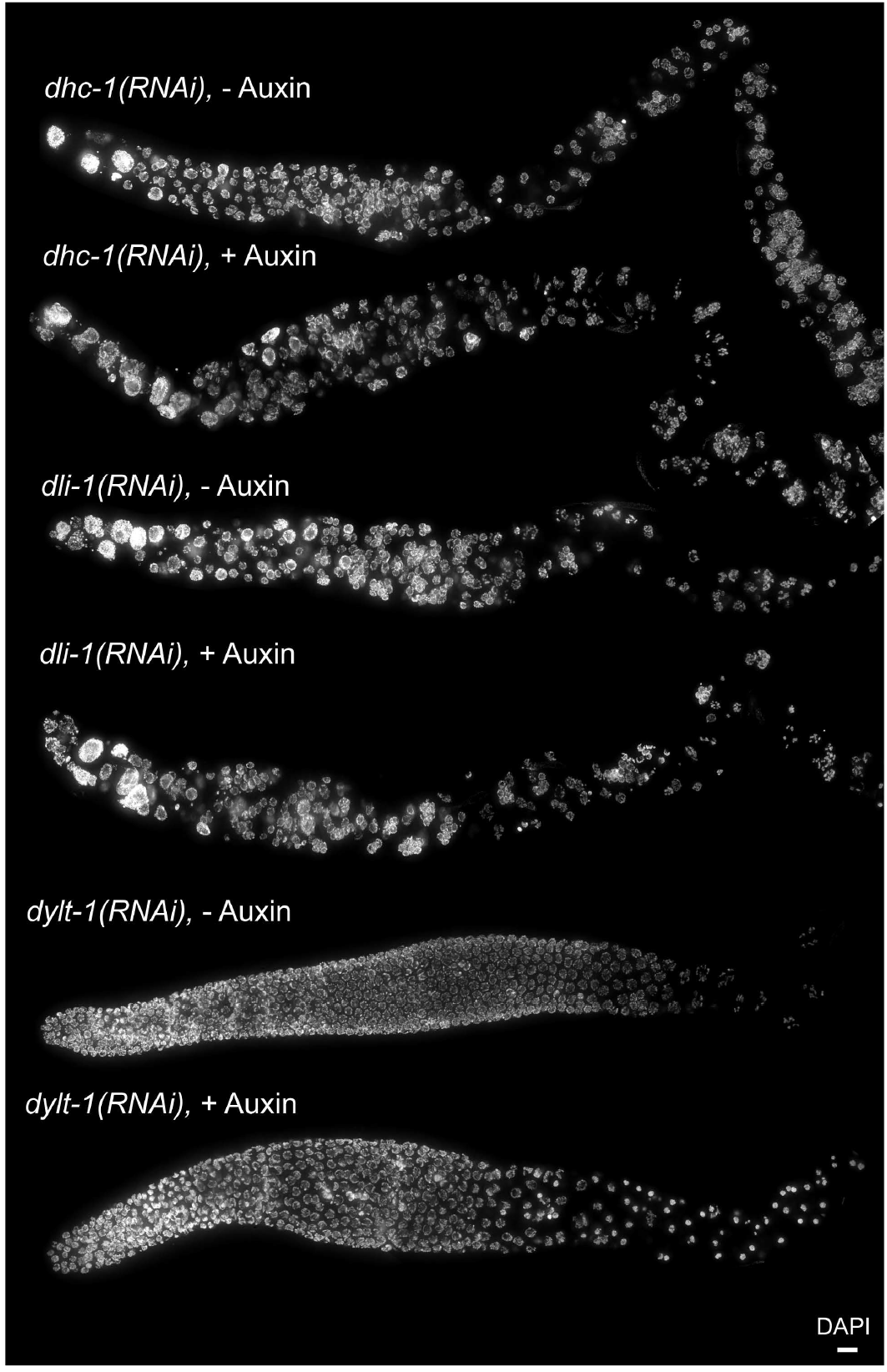
Co-depletion of DHC-1 or DLI-1, but not DYLT-1, also rescues nuclear collapse despite marked nuclear mispositioning. Nuclear morphology upon depleting DHC-1, DLI-1 or DYLT-1 in *lmn-1::AID::V5* worms ±auxin treatment. Mitotic defects are seen in the proliferative region, and meiotic nuclei are mispositioned throughout the gonad following depletion of DHC-1 or DLI-1. Scale bar, 10 µm.

**Supplementary Figure 10 (Related to Figure 3).**
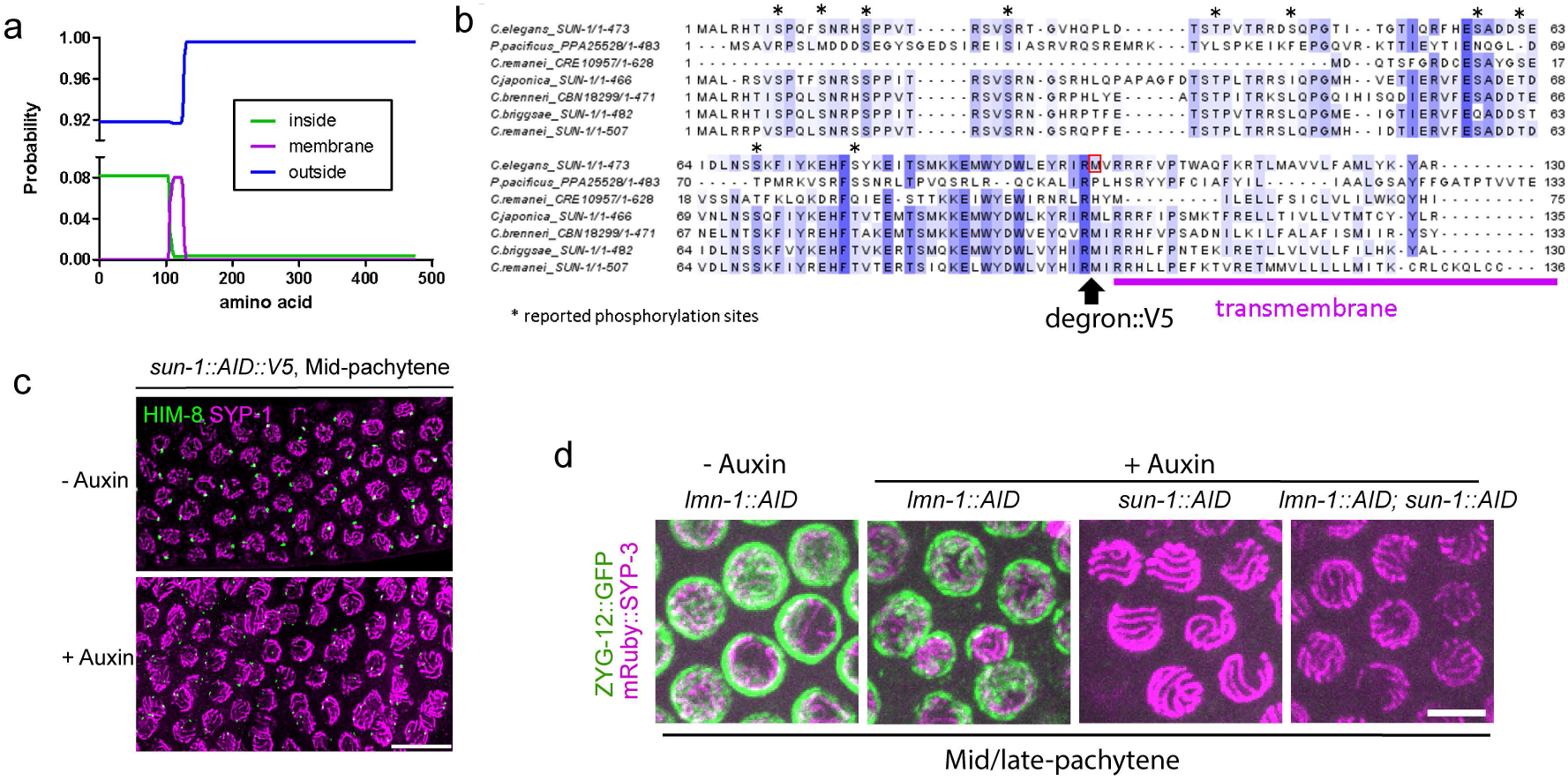
CRISPR tagging and auxin-inducible degradation of SUN-1 in *C. elegans*. **a.** Prediction of transmembrane region in *C. elegans* SUN-1 using TMHMM (http://www.cbs.dtu.dk/services/TMHMM/). Probability of amino acids inside the INM (green), outside the ONM (blue) and in the transmembrane region (magenta) is plotted. **b.** Multiple sequence alignments of nematode SUN-1 proteins were generated using Clustal Omega and visualized using Jalview, showing position of degron/V5 insertion in *C. elegans* SUN-1. **c.** X-chromosome pairing (HIM-8) and SC assembly (SYP-1) in *sun-1::AID::V5* worms without or with auxin treatment. Scale bar, 10 µm. **d.** SUN-1 is required for ZYG-12 localization at the NE of meiotic cells. Composite images showing live meiotic nuclei from mid/late pachytene in worms of indicated genotypes or treatments. ZYG-12::GFP in green and mRuby::SYP-3 in magenta. All images are maximum-intensity projections and scaled identically. Scale bar, 5 µm.

**Supplementary Figure 11 (Related to Figure 3 and 4).**
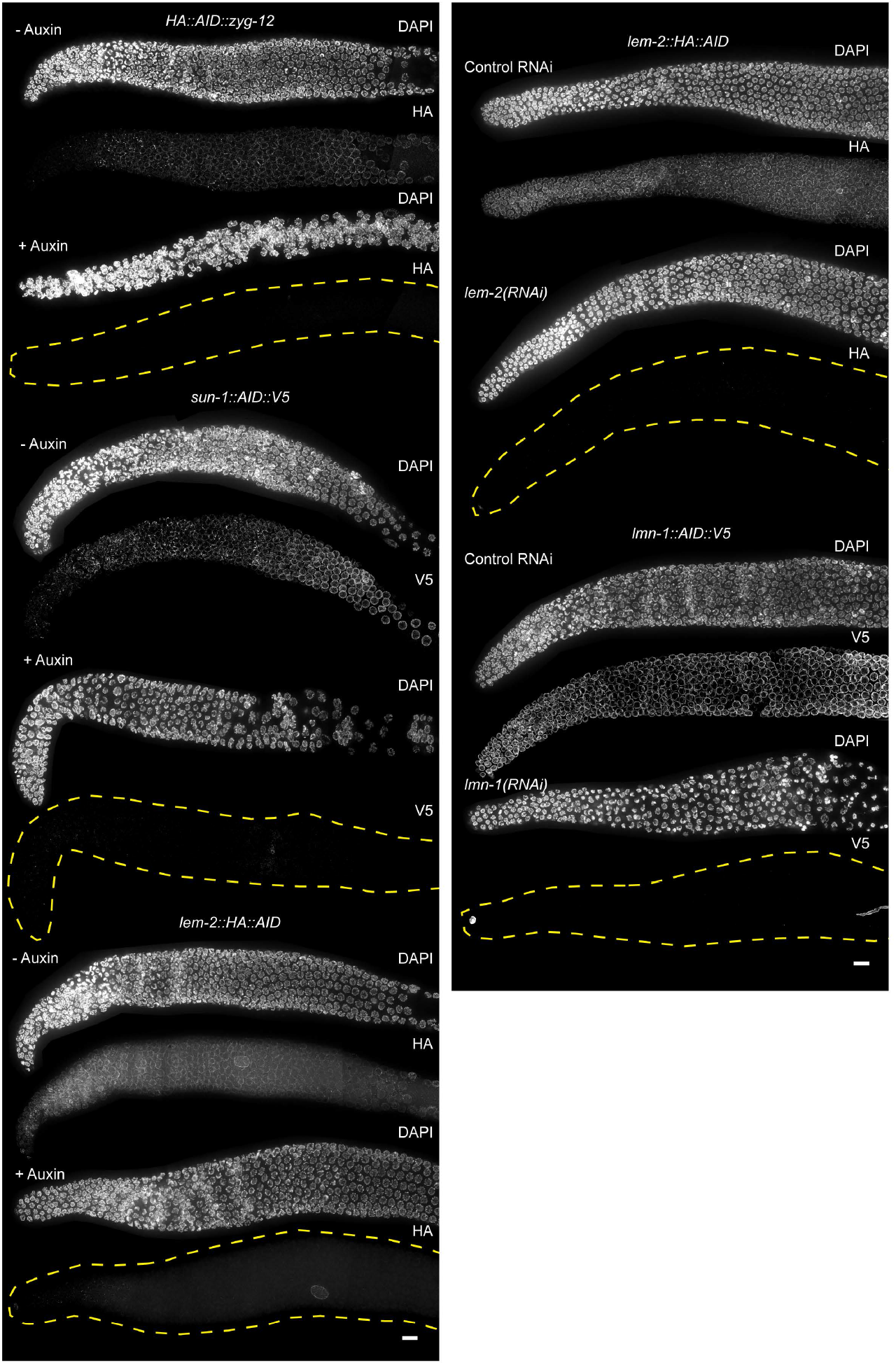
Efficacy of auxin-induced degradation or RNAi. Nuclear morphology in hermaphrodites depleted of ZYG-12, SUN-1 or LEM-2 with auxin-induced degradation, or depleted of LEM-2 or LMN-1 using RNAi. The duration of auxin treatment was at least 8 hours and the duration of feeding RNAi was 48hr. Scale bars, 10 µm.

**Supplementary Figure 12 (Related to Figure 4.**
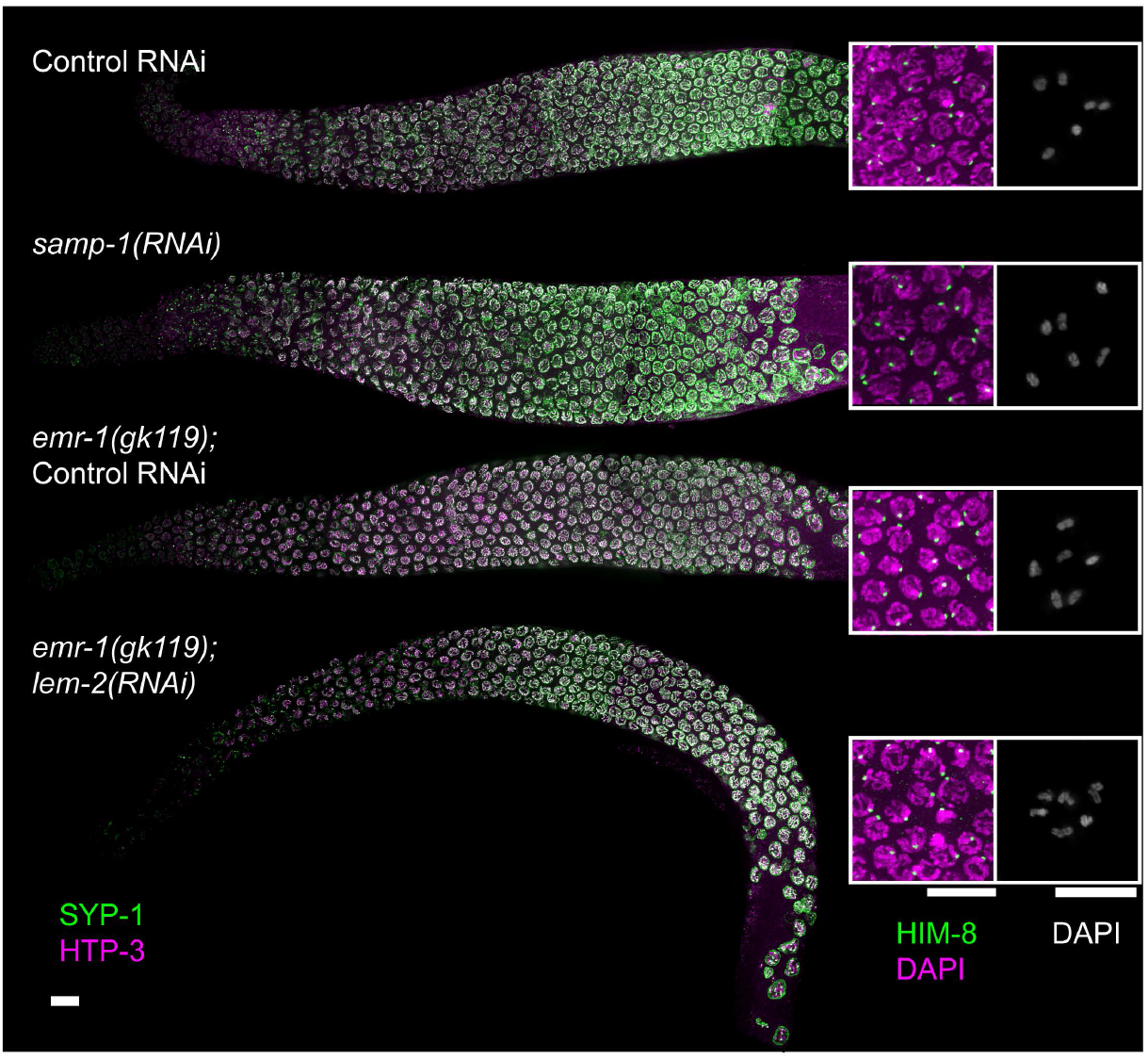
Pairing and synapsis upon depleting SAMP-1, EMR-1 or LEM-2. X-chromosome pairing (HIM-8) and SC assembly (SYP-1 and HTP-3) upon RNAi-mediated depletion of SAMP-1 or of LEM-2 in *emr-1(gk119)* mutants. Insets showing X-chromosome pairing in early pachytene and bivalents formation in diakinesis under each condition. Scale bars, 10 µm.

**Supplementary Figure 13 (Related 1to Figure 4).**
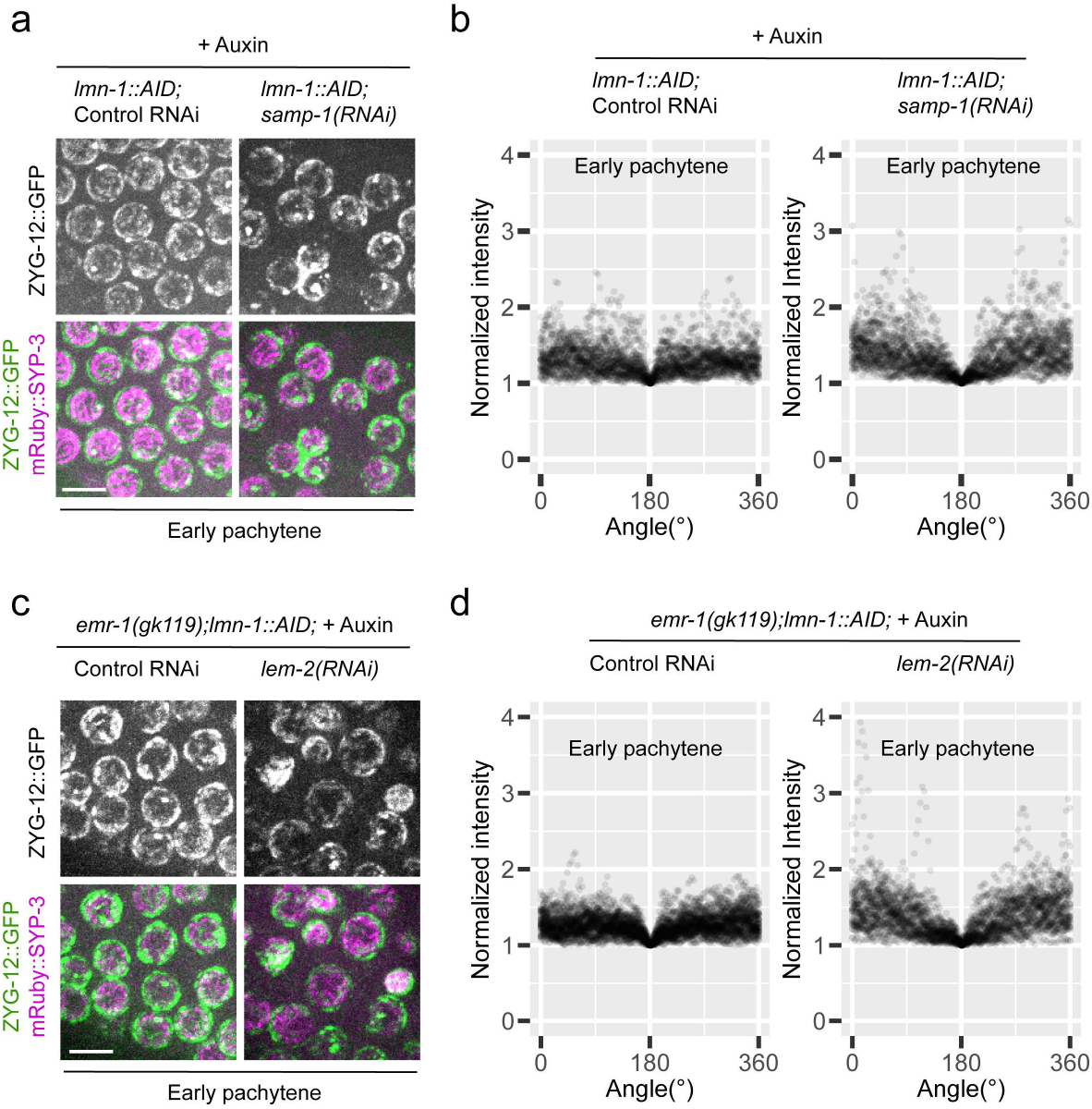
Asymmetric distribution of ZYG-12::GFP at early pachytene NE upon co-depleting LMN-1 and SAMP-1 or EMR-1/LEM-2. **a** and **c**. Composite images showing mRuby::SYP-3 (magenta) and ZYG-12::GFP (green) at the NE of early pachytene nuclei of indicated genotypes or treatments. All images are maximum-intensity projections and scaled identically per experiment. Scale bar, 5 µm. **b** and **d**. Quantification of ZYG-12::GFP fluorescence as a function of angle along the circumference of NE in early pachytene nuclei. Intensity measurement was performed as in **Supplementary Figure 8**. Because of the multiple bright foci/patches of LINC complexes at the NE in each early pachytene nucleus, data had to be aligned and normalized differently than that in Figure 3b: data were aligned and mapped so that 180° corresponds to the coordinate along the NE’s circumference with the minimum-intensity of ZYG-12::GFP, which was normalized as one. In (**b**), N = 32 early pachytene nuclei pooled from four animals were measured for each condition. *p* < 2.2e-16 between the control RNAi and *samp-1(RNAi)* with 2-way ANOVA. In (**d**), N = 51 (Control RNAi) and 35 (*lem-2(RNAi)*) early pachytene nuclei pooled from seven or four animals were measured. *p* < 2.2e-16 between the control RNAi and *lem-2(RNAi)* with 2-way ANOVA. See **Statistical Source Data**.

## SUPPLEMENTARY VIDEOS

**Supplementary Video 1.** Two representative time-lapse recordings of SUN-1::mRuby at the NE of meiotic nuclei in late meiotic prophase. Arrow points to a collapsing nucleus. Note many meiotic nuclei with asymmetrically distributed SUN-1::mRuby at the NE. Time stamp is hr:min:sec. Scale bars, 10 µm.

**Supplementary Video 2.** An example of drift-corrected time-lapse recordings of SUN-1::mRuby patch movement on the NE of transition zone nuclei, using reference frame followed by 3D particle tracking in Imaris. Time stamp is hr:min:sec. Scale bar, 5 µm.

**Supplementary Video 3.** Side-by-side comparison of time-lapse recordings of the movement of SUN-1::mRuby patches or foci during different stages of meiosis, with or without LMN-1 depletion. Time stamp is hr:min:sec. Scale bars, 5 µm.

**Supplementary Video 4.** Time-lapse recording of a dual-color labeled oocyte nucleus during diplotene collapse. Green, ZYG-12::GFP; Magenta, mRuby::SYP-3. Scale bar, 5 µm.

**Supplementary Video 5.** Time-lapse recording of another dual-color labeled diplotene nucleus during collapse. The frame showing initial contact between NE and SC is annotated. Green, ZYG-12::GFP; Magenta, mRuby::SYP-3. Time stamp is hr:min:sec. Scale bar, 2 µm.

## SUPPLEMENTARY TABLES

**Supplementary Table 1.** Quantification of brood size, embryo viability, and male self-progeny of worm strains with new AID alleles generated in this study (*lmn-1::AID::V5, HA::AID::zyg-12, sun-1::AID::V5, lem-2::AID::HA*). Sample size n indicates the number of hermaphrodites examined. Note that high levels of uncertainty are usually associated with counting deformed eggs resulting from auxin treatment.

**Supplementary Table 2.** New alleles generated in this study.

**Supplementary Table 3.** Genotypes of worm strains generated and used in this study.

**Supplementary Table 4.** Sequences of gRNAs, repair templates, and DNA primers used to genotype edited progeny.

**Supplementary Table 5.** RNAi clones used in this study.

## SOURCE DATA: STATISTICAL SOURCE DATA

Additional output of statistical tests performed in this study (with related Figure or Supplementary Figure numbers).

## Notes

### Competing Interest Statement

The authors have declared no competing interest.

