## Supplementary Statistics Data for "A cooperative network at the nuclear envelope counteracts LINC-mediated forces during oogenesis in *C. elegans*"

### SOURCE DATA

#### STATISTICAL SOURCE DATA

**TABLE OF CONTENT:**

Additional output of statistical analyses for data presented in main figures and Supplementary figures are as following (statistics of interests are highlighted):

| Output of statistical analyses | Figure(s) |
| --- | --- |
| Two-way ANOVA on the mean and relative standard deviation of SUN-1::mRuby intensity; | Figure 2b,c |
| Statistics for asymmetry of diplotene LINC distribution, re-binned, t-test for peak intensity value between 170-190° | Figure 3b |
| One-way ANOVA and post hoc pairwise t-tests for comparing normalized nuclear sizes | Figure 3e |
| For comparing the percentage of nuclear collapse in each zone, post hoc pair-wise comparison of proportions | Figure 4b |
| For comparing the percentage of nuclear collapse in each zone, post-hoc pair-wise comparison of proportions | Figure 4d |
| Unpaired two-sample two-sided t-test for comparing normalized nuclear radius, nuclear volume (derived), DAPI volume as well as DAPI integrated intensity in control or Auxin treated animals | Supplementary Figure 3c |
| For comparing the proportion of nuclei with at least one RAD-51 focus in each zone across groups, post hoc pair wise comparison of proportions | Supplementary Figure 4c |
| For comparing the percentage of X-chromosome pairing or complete synapsis | Supplementary Figure 6c |
| Statistics for comparing the extent of asymmetry | Supplementary Figure 13b |
| Statistics for comparing the extent of asymmetry | Supplementary Figure 13d |

**Two-way ANOVA on the mean and relative standard deviation of SUN-1::mRuby intensity; related to Figure 2b,c:**

```
> res.aov3 <- aov(mean ~ zone * group, data = df_norm)
> summary(res.aov3)
             Df Sum Sq Mean Sq F value Pr(>F)    
zone          3  98.51   32.84  101.35 <2e-16 ***
group         1  96.22   96.22  296.99 <2e-16 ***
zone:group    3  46.10   15.37   47.44 <2e-16 ***
Residuals   232  75.16    0.32                   
---
Signif. codes:  0 ‘***’ 0.001 ‘**’ 0.01 ‘*’ 0.05 ‘.’ 0.1 ‘ ’ 1
> res.aov4 <- aov(coef_v ~ zone * group, data = df_norm)
> summary(res.aov4)
             Df Sum Sq Mean Sq F value   Pr(>F)    
zone          3 0.7650  0.2550  55.452  < 2e-16 ***
group         1 0.5890  0.5890 128.098  < 2e-16 ***
zone:group    3 0.1252  0.0417   9.073 1.05e-05 ***
Residuals   232 1.0668  0.0046                     
---
Signif. codes:  0 ‘***’ 0.001 ‘**’ 0.01 ‘*’ 0.05 ‘.’ 0.1 ‘ ’ 1
```

**Statistics for asymmetry of diplotene LINC distribution, re-binned, t-test for peak intensity value between 170-190°; related to Figure 3b:**

**L4440 -/+Aux (1 being -Aux, 2 being + Aux):**

```
> df_t <- rbind(df_t1,df_t2)
> df_t$id <- factor(df_t$id)
> res.ftest <- var.test(Y ~ id, data = df_t)
> res.ftest
F test to compare two variances
data:  Y by id
F = 0.056958, num df = 420, denom df = 295, p-value < 2.2e-16
alternative hypothesis: true ratio of variances is not equal to 1
95 percent confidence interval:
 0.04603585 0.07019336
sample estimates:
ratio of variances
         0.0569576
>
> t.test(Y~id, data = df_t, alternative = "two.sided", var.equal = TRUE)
Two Sample t-test
data:  Y by id
t = -27.112, df = 715, p-value < 2.2e-16
alternative hypothesis: true difference in means is not equal to 0
95 percent confidence interval:
 -0.8531666 -0.7379485
sample estimates:
mean in group 1 mean in group 2
       1.197693        1.993250
> t.test(Y~id, data = df_t, alternative = "two.sided", var.equal = FALSE)
Welch Two Sample t-test
data:  Y by id
t = -23.171, df = 318.74, p-value < 2.2e-16
alternative hypothesis: true difference in means is not equal to 0
95 percent confidence interval:
 -0.8631076 -0.7280075
sample estimates:
mean in group 1 mean in group 2
       1.197693        1.993250
```

100*(1.993250-1.197693)/1.197693 = 66.424117 percent difference;

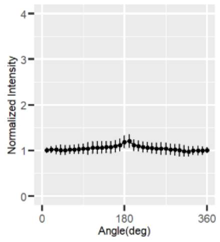


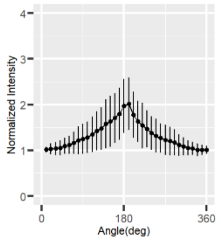


**Dnc-1 -/+ Aux (1 being -Aux, 2 being +Aux):**

```
> df_t <- rbind(df_t1,df_t2)
> df_t$id <- factor(df_t$id)
> res.ftest <- var.test(Y ~ id, data = df_t)
> res.ftest
F test to compare two variances
data:  Y by id
F = 0.84728, num df = 326, denom df = 411, p-value = 0.1166
alternative hypothesis: true ratio of variances is not equal to 1
95 percent confidence interval:
 0.6904682 1.0422827
sample estimates:
ratio of variances
         0.8472795
```

```
>
> t.test(Y~id, data = df_t, alternative = "two.sided", var.equal = TRUE)
Two Sample t-test
data:  Y by id
t = 1.1265, df = 737, p-value = 0.2603
alternative hypothesis: true difference in means is not equal to 0
95 percent confidence interval:
 -0.01169140  0.04317295
sample estimates:
mean in group 1 mean in group 2
       1.253348        1.237607
100*(1.237607-1.253348)/1.253348 = -1.255916 percent different;

> t.test(Y~id, data = df_t, alternative = "two.sided", var.equal = FALSE)
Welch Two Sample t-test
data:  Y by id
t = 1.1373, df = 721, p-value = 0.2558
alternative hypothesis: true difference in means is not equal to 0
95 percent confidence interval:
 -0.01143242  0.04291397
sample estimates:
mean in group 1 mean in group 2
       1.253348        1.237607
```

**Dlc-1 -/+ Aux (1 being -Aux, 2 being +Aux):**

```
> df_t <- rbind(df_t1,df_t2)
> df_t$id <- factor(df_t$id)
> res.ftest <- var.test(Y ~ id, data = df_t)
> res.ftest
F test to compare two variances
data:  Y by id
F = 0.71667, num df = 465, denom df = 436, p-value = 0.0004144
alternative hypothesis: true ratio of variances is not equal to 1
95 percent confidence interval:
 0.5954561 0.8620879
sample estimates:
ratio of variances
         0.7166744
>
> t.test(Y~id, data = df_t, alternative = "two.sided", var.equal = TRUE)
Two Sample t-test
data:  Y by id
t = -4.748, df = 901, p-value = 2.388e-06
alternative hypothesis: true difference in means is not equal to 0
95 percent confidence interval:
 -0.06275017 -0.02604620
sample estimates:
mean in group 1 mean in group 2
       1.162345        1.206744
> t.test(Y~id, data = df_t, alternative = "two.sided", var.equal = FALSE)
Welch Two Sample t-test
data:  Y by id
t = -4.7229, df = 856.32, p-value = 2.716e-06
alternative hypothesis: true difference in means is not equal to 0
95 percent confidence interval:
 -0.06284911 -0.02594727
sample estimates:
mean in group 1 mean in group 2
       1.162345        1.206744

100*(1.206744-1.162345)/1.152345 = 3.852926 percent different;
```

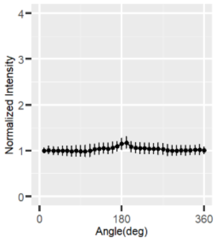
Normalized Intensity
4
3
2
1
0
0
180
360
Angle(deg)
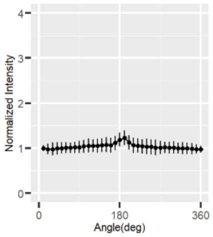
Normalized Intensity
4
3
2
1
0
0
180
360
Angle(deg)

**One-way ANOVA and post hoc pairwise t-tests for comparing normalized nuclear sizes; related to Figure 3e:**

```
> res.aov <- aov(volume~group, data = data_t1)
> summary(res.aov)
              Df Sum Sq Mean Sq F value Pr(>F)    
group         17  447.5  26.326   162.9 <2e-16 ***
Residuals   1945  314.2   0.162                   
---
Signif. codes:  0 ‘***’ 0.001 ‘**’ 0.01 ‘*’ 0.05 ‘.’ 0.1 ‘ ’ 1
> pairwise.t.test(data_t1$volume, data_t1$group,
+                 p.adjust.method = "BH")
Pairwise comparisons using t tests with pooled SD 
data:  data_t1$volume and data_t1$group 
                      - Auxin LMN-1 AID SUN-1 AID LMN-1, SUN-1 AID ZYG-12 AID LMN-1, ZYG-12 AID L4440  
LMN-1 AID             < 2e-16 -         -         -                -          -                 -      
SUN-1 AID             0.05963 < 2e-16   -         -                -          -                 -      
LMN-1, SUN-1 AID      0.03323 < 2e-16   0.76377   -                -          -                 -      
ZYG-12 AID            2.2e-08 < 2e-16   3.3e-15   3.1e-15          -          -                 -      
LMN-1, ZYG-12 AID     5.2e-06 < 2e-16   1.6e-11   1.1e-11          0.40562    -                 -      
L4440                 8.4e-09 < 2e-16   3.3e-05   0.00022          < 2e-16    < 2e-16           -      
L4440, LMN-1 AID      < 2e-16 0.30272   < 2e-16   < 2e-16          < 2e-16    < 2e-16           < 2e-16
dnc-1 RNAi            4.0e-05 < 2e-16   0.01386   0.03696          < 2e-16    < 2e-16           0.18167
dnc-1 RNAi, LMN-1 AID 0.00195 < 2e-16   0.18419   0.32447          < 2e-16    3.0e-14           0.00969
lis-1 RNAi            2.5e-15 < 2e-16   1.9e-10   6.3e-09          < 2e-16    < 2e-16           0.08788
lis-1 RNAi, LMN-1 AID 0.09022 < 2e-16   0.85319   0.62489          1.1e-14    4.7e-11           1.3e-05
dlc-1 RNAi            1.7e-05 < 2e-16   0.00870   0.02535          < 2e-16    < 2e-16           0.20281
dlc-1 RNAi, LMN-1 AID 0.00035 < 2e-16   0.07425   0.16354          < 2e-16    4.0e-16           0.02441
emr-1                 0.20498 < 2e-16   0.76336   0.58174          2.5e-10    6.3e-08           0.00010
emr-1, LMN-1 AID      < 2e-16 0.63772   < 2e-16   < 2e-16          < 2e-16    < 2e-16           < 2e-16
LEM-2 AID             9.8e-06 < 2e-16   0.00685   0.02142          < 2e-16    < 2e-16           0.18975
LEM-2, LMN-1 AID      < 2e-16 0.97147   < 2e-16   < 2e-16          < 2e-16    < 2e-16           < 2e-16
                      L4440, LMN-1 AID dnc-1 RNAi dnc-1 RNAi, LMN-1 AID lis-1 RNAi lis-1 RNAi, LMN-1 AID dlc-1 
RNAi   
LMN-1 AID             -                -          -                     -          -                     -      
SUN-1 AID             -                -          -                     -          -                     -      
LMN-1, SUN-1 AID      -                -          -                     -          -                     -      
ZYG-12 AID            -                -          -                     -          -                     -      
LMN-1, ZYG-12 AID     -                -          -                     -          -                     -      
L4440                 -                -          -                     -          -                     -      
L4440, LMN-1 AID      -                -          -                     -          -                     -      
dnc-1 RNAi            < 2e-16          -          -                     -          -                     -      
dnc-1 RNAi, LMN-1 AID < 2e-16          0.28264    -                     -          -                     -      
lis-1 RNAi            < 2e-16          0.00166    4.9e-06               -          -                     -      
lis-1 RNAi, LMN-1 AID < 2e-16          0.00790    0.12534               3.9e-11    -                     -      
dlc-1 RNAi            < 2e-16          0.93586    0.23208               0.00187    0.00468               -      
dlc-1 RNAi, LMN-1 AID < 2e-16          0.46564    0.72562               2.0e-05    0.04520               0.39882
emr-1                 < 2e-16          0.01473    0.14955               8.9e-09    0.89237               0.01005
emr-1, LMN-1 AID      0.63772          < 2e-16    < 2e-16               < 2e-16    < 2e-16               < 2e-16
LEM-2 AID             < 2e-16          0.93586    0.22034               0.00135    0.00346               0.99743
LEM-2, LMN-1 AID      0.24866          < 2e-16    < 2e-16               < 2e-16    < 2e-16               < 2e-16
                      dlc-1 RNAi, LMN-1 AID emr-1   emr-1, LMN-1 AID LEM-2 AID
LMN-1 AID             -                     -       -                -        
SUN-1 AID             -                     -       -                -        
LMN-1, SUN-1 AID      -                     -       -                -        
ZYG-12 AID            -                     -       -                -        
LMN-1, ZYG-12 AID     -                     -       -                -        
L4440                 -                     -       -                -        
L4440, LMN-1 AID      -                     -       -                -        
dnc-1 RNAi            -                     -       -                -        
dnc-1 RNAi, LMN-1 AID -                     -       -                -        
lis-1 RNAi            -                     -       -                -        
lis-1 RNAi, LMN-1 AID -                     -       -                -        
dlc-1 RNAi            -                     -       -                -        
dlc-1 RNAi, LMN-1 AID -                     -       -                -        
emr-1                 0.06691               -       -                -        
emr-1, LMN-1 AID      < 2e-16               < 2e-16 -                -        
LEM-2 AID             0.38777               0.00856 < 2e-16          -        
LEM-2, LMN-1 AID      < 2e-16               < 2e-16 0.59395          < 2e-16  
P value adjustment method: BH
```

**For comparing the percentage of nuclear collapse in each zone, post-hoc pair-wise comparison of proportions; related to Figure 4b:**

#samp-1 zone 1~6 (sequential sections below); "1" ~ "5" in each section are Control RNAi - Aux, Control RNAi + Aux, samp-1(RNAi) - Aux, samp-1(RNAi) + Aux, sun-1::AID and samp-1(RNAi) + Aux

```
> prop.test(x=c(5,3,4, 16, 7),n=c(281, 299, 261, 259,  349))
5-sample test for equality of proportions without continuity correction
data:  c(5, 3, 4, 16, 7) out of c(281, 299, 261, 259, 349)
X-squared = 19.676, df = 4, p-value = 0.0005787
alternative hypothesis: two.sided
sample estimates:
    prop 1     prop 2     prop 3     prop 4     prop 5
0.01779359 0.01003344 0.01532567 0.06177606 0.02005731
> pairwise.prop.test(x=c(5,3,4, 16, 7),n=c(281, 299, 261, 259, 349),p.adjust.method =
"BH")
Pairwise comparisons using Pairwise comparison of proportions
data:  c(5, 3, 4, 16, 7) out of c(281, 299, 261, 259, 349)
  1     2     3     4
2 1.000 -     -     -
3 1.000 1.000 -     -
4 0.039 0.018 0.039 -
5 1.000 0.953 1.000 0.039
P value adjustment method: BH


> prop.test(x=c(4, 1, 3, 63, 10),n=c(316, 306, 379, 305, 433))
5-sample test for equality of proportions without continuity correction
data:  c(4, 1, 3, 63, 10) out of c(316, 306, 379, 305, 433)
X-squared = 215.02, df = 4, p-value < 2.2e-16
alternative hypothesis: two.sided
sample estimates:
     prop 1      prop 2      prop 3      prop 4      prop 5
0.012658228 0.003267974 0.007915567 0.206557377 0.023094688
> pairwise.prop.test(x=c(4, 1, 3, 63, 10),n=c(316, 306, 379, 305, 433),p.adjust.method =
"BH")
Pairwise comparisons using Pairwise comparison of proportions
data:  c(4, 1, 3, 63, 10) out of c(316, 306, 379, 305, 433)
  1       2       3       4
2 0.55    -       -       -
3 0.81    0.81    -       -
4 4.8e-14 2.3e-15 < 2e-16 -
5 0.55    0.12    0.25    2.3e-15
P value adjustment method: BH


> prop.test(x=c(6,64, 20, 175,15),n=c(405, 389, 459, 365, 429))
5-sample test for equality of proportions without continuity correction
data:  c(6, 64, 20, 175, 15) out of c(405, 389, 459, 365, 429)
X-squared = 487.98, df = 4, p-value < 2.2e-16
alternative hypothesis: two.sided
sample estimates:
    prop 1     prop 2     prop 3     prop 4     prop 5
0.01481481 0.16452442 0.04357298 0.47945205 0.03496503
> pairwise.prop.test(x=c(6,64, 20, 175,15),n=c(405, 389, 459, 365, 429),p.adjust.method =
"BH")
Pairwise comparisons using Pairwise comparison of proportions
data:  c(6, 64, 20, 175, 15) out of c(405, 389, 459, 365, 429)
  1       2       3       4
2 5.2e-13 -       -       -
3 0.029   1.2e-08 -       -
4 < 2e-16 < 2e-16 < 2e-16 -
5 0.113   1.3e-09 0.627   < 2e-16
P value adjustment method: BH



> prop.test(x=c(12, 50, 10, 82, 10),n=c(326, 287, 407, 195, 354))
5-sample test for equality of proportions without continuity correction
```

```
data:  c(12, 50, 10, 82, 10) out of c(326, 287, 407, 195, 354)
X-squared = 288.68, df = 4, p-value < 2.2e-16
alternative hypothesis: two.sided
sample estimates:
    prop 1     prop 2     prop 3     prop 4     prop 5 
0.03680982 0.17421603 0.02457002 0.42051282 0.02824859 
> pairwise.prop.test(x=c(12, 50, 10, 82, 10),n=c(326, 287, 407, 195, 354),p.adjust.method
= "BH")
Pairwise comparisons using Pairwise comparison of proportions 
data:  c(12, 50, 10, 82, 10) out of c(326, 287, 407, 195, 354) 
  1       2       3       4      
2 5.6e-08 -       -       -      
3 0.57    3.2e-11 -       -      
4 < 2e-16 8.3e-09 < 2e-16 -      
5 0.75    1.3e-09 0.93    < 2e-16
P value adjustment method: BH 


> prop.test(x=c(0, 108, 3, 151, 8),n=c(50, 147, 145, 162, 243))
5-sample test for equality of proportions without continuity correction
data:  c(0, 108, 3, 151, 8) out of c(50, 147, 145, 162, 243)
X-squared = 532.18, df = 4, p-value < 2.2e-16
alternative hypothesis: two.sided
sample estimates:
    prop 1     prop 2     prop 3     prop 4     prop 5 
0.00000000 0.73469388 0.02068966 0.93209877 0.03292181 
> pairwise.prop.test(x=c(0, 108, 3, 151, 8),n=c(50, 147, 145, 162, 243),p.adjust.method =
"BH")
Pairwise comparisons using Pairwise comparison of proportions 
data:  c(0, 108, 3, 151, 8) out of c(50, 147, 145, 162, 243) 
  1       2       3       4      
2 < 2e-16 -       -       -      
3 0.72    < 2e-16 -       -      
4 < 2e-16 7.6e-06 < 2e-16 -      
5 0.51    < 2e-16 0.72    < 2e-16
P value adjustment method: BH 


> prop.test(x=c(0, 91, 0, 103, 6),n=c(18, 92, 19, 106, 132))
5-sample test for equality of proportions without continuity correction
data:  c(0, 91, 0, 103, 6) out of c(18, 92, 19, 106, 132)
X-squared = 328.16, df = 4, p-value < 2.2e-16
alternative hypothesis: two.sided
sample estimates:
    prop 1     prop 2     prop 3     prop 4     prop 5 
0.00000000 0.98913043 0.00000000 0.97169811 0.04545455 
> pairwise.prop.test(x=c(0, 91, 0, 103, 6),n=c(18, 92, 19, 106, 132),p.adjust.method =
"BH")
Pairwise comparisons using Pairwise comparison of proportions 
data:  c(0, 91, 0, 103, 6) out of c(18, 92, 19, 106, 132) 
  1      2      3      4     
2 <2e-16 -      -      -     
3 -      <2e-16 -      -     
4 <2e-16 0.78   <2e-16 -     
5 0.78   <2e-16 0.78   <2e-16
P value adjustment method: BH 
```

**For comparing the percentage of nuclear collapse in each zone, post hoc pair-wise comparison of proportions; related to Figure 4d:**

#Lem-2, zone 1~6 (sequential sections below); “1” ~ “5” in each section are Control RNAi - Aux, Control RNAi + Aux, lem-2(RNAi) - Aux, lem-2(RNAi) + Aux, sun-1::AID and lem-2(RNAi) + Aux

```
> prop.test(x=c(5,4,92,114,83),n=c(258,287,244,188,201),correct = TRUE)
5-sample test for equality of proportions without continuity correction
data:  c(5, 4, 92, 114, 83) out of c(258, 287, 244, 188, 201)
X-squared = 332.62, df = 4, p-value < 2.2e-16
alternative hypothesis: two.sided
sample estimates:
    prop 1     prop 2     prop 3     prop 4     prop 5
0.01937984 0.01393728 0.37704918 0.60638298 0.41293532
> pairwise.prop.test(x=c(5,4,92,114,83),n=c(258,287,244,188,201),p.adjust.method = "BH")
Pairwise comparisons using Pairwise comparison of proportions
data:  c(5, 4, 92, 114, 83) out of c(258, 287, 244, 188, 201)
  1       2       3       4
2 0.87194 -       -       -
3 < 2e-16 < 2e-16 -       -
4 < 2e-16 < 2e-16 5.1e-06 -
5 < 2e-16 < 2e-16 0.55608 0.00026
P value adjustment method: BH

> prop.test(x=c(9,5,42,104,14),n=c(342, 320, 376,  208, 241))
5-sample test for equality of proportions without continuity correction
data:  c(9, 5, 42, 104, 14) out of c(342, 320, 376, 208, 241)
X-squared = 362.55, df = 4, p-value < 2.2e-16
alternative hypothesis: two.sided
sample estimates:
    prop 1     prop 2     prop 3     prop 4     prop 5
0.02631579 0.01562500 0.11170213 0.50000000 0.05809129
> pairwise.prop.test(x=c(9,5,42,104,14),n=c(342, 320,376, 208,241),p.adjust.method =
"BH")
Pairwise comparisons using Pairwise comparison of proportions
data:  c(9, 5, 42, 104, 14) out of c(342, 320, 376, 208, 241)
  1       2       3       4
2 0.493   -       -       -
3 2.8e-05 2.1e-06 -       -
4 < 2e-16 < 2e-16 < 2e-16 -
5 0.094   0.017   0.043   < 2e-16
P value adjustment method: BH


> prop.test(x=c(6,82,26, 163,9),n=c(448,373,439,235, 231))
5-sample test for equality of proportions without continuity correction
data:  c(6, 82, 26, 163, 9) out of c(448, 373, 439, 235, 231)
X-squared = 619.68, df = 4, p-value < 2.2e-16
alternative hypothesis: two.sided
sample estimates:
    prop 1     prop 2     prop 3     prop 4     prop 5
0.01339286 0.21983914 0.05922551 0.69361702 0.03896104
> pairwise.prop.test(x=c(6,82,26,163,9),n=c(448, 373, 439, 235,231),p.adjust.method =
"BH")
Pairwise comparisons using Pairwise comparison of proportions
data:  c(6, 82, 26, 163, 9) out of c(448, 373, 439, 235, 231)
  1       2       3       4
2 < 2e-16 -       -       -
3 0.00063 6.3e-11 -       -
4 < 2e-16 < 2e-16 < 2e-16 -
5 0.06800 4.5e-09 0.34835 < 2e-16
P value adjustment method: BH


> prop.test(x=c(19,  91,  25,  159, 13),n=c(437, 292,318,189,127))
5-sample test for equality of proportions without continuity correction
data:  c(19, 91, 25, 159, 13) out of c(437, 292, 318, 189, 127)
X-squared = 556.4, df = 4, p-value < 2.2e-16
alternative hypothesis: two.sided
sample estimates:
```

```
    prop 1     prop 2     prop 3     prop 4     prop 5 
0.04347826 0.31164384 0.07861635 0.84126984 0.10236220 
> pairwise.prop.test(x=c(19,91,25,159,13),n=c(437,292,318,189,127),p.adjust.method = 
"BH")
Pairwise comparisons using Pairwise comparison of proportions 
data:  c(19, 91, 25, 159, 13) out of c(437, 292, 318, 189, 127) 
  1       2       3       4      
2 < 2e-16 -       -       -      
3 0.067   8.5e-13 -       -      
4 < 2e-16 < 2e-16 < 2e-16 -      
5 0.026   1.3e-05 0.534   < 2e-16
P value adjustment method: BH 

> prop.test(x=c(12,154,8,66,13),n=c(198, 197, 94, 76, 87))
5-sample test for equality of proportions without continuity correction
data:  c(12, 154, 8, 66, 13) out of c(198, 197, 94, 76, 87)
X-squared = 349.02, df = 4, p-value < 2.2e-16
alternative hypothesis: two.sided
sample estimates:
    prop 1     prop 2     prop 3     prop 4     prop 5 
0.06060606 0.78172589 0.08510638 0.86842105 0.14942529 
> pairwise.prop.test(x=c(12,154,8,66,13),n=c(198,197,94, 76, 87),p.adjust.method = "BH")
Pairwise comparisons using Pairwise comparison of proportions 
data:  c(12, 154, 8, 66, 13) out of c(198, 197, 94, 76, 87) 
  1      2      3      4     
2 <2e-16 -      -      -     
3 0.599  <2e-16 -      -     
4 <2e-16 0.183  <2e-16 -     
5 0.038  <2e-16 0.293  <2e-16
P value adjustment method: BH 


> prop.test(x=c(1,80, 5,41,2),n=c(24, 86,  26, 43, 39))
5-sample test for equality of proportions without continuity correction
data:  c(1, 80, 5, 41, 2) out of c(24, 86, 26, 43, 39)
X-squared = 158.47, df = 4, p-value < 2.2e-16
alternative hypothesis: two.sided
sample estimates:
    prop 1     prop 2     prop 3     prop 4     prop 5 
0.04166667 0.93023256 0.19230769 0.95348837 0.05128205 
> pairwise.prop.test(x=c(1,80,5,41,2),n=c(24,86,26, 43, 39),p.adjust.method = "BH")
Pairwise comparisons using Pairwise comparison of proportions 
data:  c(1, 80, 5, 41, 2) out of c(24, 86, 26, 43, 39) 
  1       2       3       4      
2 < 2e-16 -       -       -      
3 0.29    2.4e-13 -       -      
4 1.9e-12 1.00    7.5e-10 -      
5 1.00    < 2e-16 0.24    6.3e-15
P value adjustment method: BH 
```

**Unpaired two-sample two-sided t-test for comparing normalized nuclear radius, nuclear volume (derived), DAPI volume as well as DAPI integrated intensity in control or Auxin treated animals; related to Supplementary Figure 3c:**

```
# Radius
> t.test(length~zone, data = df_norm, alternative = "two.sided", var.equal = TRUE)
Two Sample t-test
data:  length by zone
t = 11.886, df = 158, p-value < 2.2e-16
alternative hypothesis: true difference in means is not equal to 0
95 percent confidence interval:
 0.2202022 0.3079699
sample estimates:
mean in group Late Pachytene      mean in group Diplotene
                    1.000000                     0.735914

# Radius^3 is volume:
> res.ftest <- var.test(length ~ zone, data = df_norm3)
> res.ftest
F test to compare two variances
data:  length by zone
F = 1.2472, num df = 79, denom df = 79, p-value = 0.3282
alternative hypothesis: true ratio of variances is not equal to 1
95 percent confidence interval:
 0.7998712 1.9447723
sample estimates:
ratio of variances
          1.247224
> t.test(length~zone, data = df_norm3, alternative = "two.sided", var.equal = TRUE)
Two Sample t-test
data:  length by zone
t = 10.651, df = 158, p-value < 2.2e-16
alternative hypothesis: true difference in means is not equal to 0
95 percent confidence interval:
 0.4917448 0.7156346
sample estimates:
mean in group Late Pachytene      mean in group Diplotene
                     1.05272                      0.44903


# dapi volume:
> t.test(volume~zone, data = dat_norm, alternative = "two.sided", var.equal = TRUE)
Two Sample t-test
data:  volume by zone
t = 12.645, df = 124, p-value < 2.2e-16
alternative hypothesis: true difference in means is not equal to 0
95 percent confidence interval:
 0.3641613 0.4993231
sample estimates:
mean in group Late Pachytene      mean in group Diplotene
                   1.0000000                    0.5682578

# dapi integrated intensity:
> t.test(intensity~zone, data = dat_norm, alternative = "two.sided", var.equal = FALSE)
Welch Two Sample t-test
data:  intensity by zone
t = -0.5734, df = 106.76, p-value = 0.5676
alternative hypothesis: true difference in means is not equal to 0
95 percent confidence interval:
 -0.13188821  0.07271006
sample estimates:
mean in group Late Pachytene      mean in group Diplotene
                    1.000000                     1.029589

[N = 58 (LP), 68 (Dip) for volume; N = 62 (LP), 64 (Dip) for intensity; N = 80 (LP), 80 (Dip) for radius.]
```

**For comparing the proportion of nuclei with at least one RAD-51 focus in each zone across groups, post hoc pair wise comparison of proportions; related to Supplementary Figure 4c:**

```
# zone 1~5 (sequential sections below); "1" ~ "6" in each section are Control, SPO-11, LMN-1, LMN-1 and SPO-11 (double), LMN-1 and SPO-11 and SUN-1 (triple), and SUN-1:

[1] "this is for zone 1"
6-sample test for equality of proportions without continuity correction
data:  c(one_ctrl$count[k], one_spo$count[k], one_lmn$count[k], one_double$count[k],  out
of c(tot_ctrl$count[k], tot_spo$count[k], tot_lmn$count[k], tot_double$count[k],
one_triple$count[k], one_sun$count[k]) out of     tot_triple$count[k], tot_sun$count[k])
X-squared = 82.788, df = 5, p-value < 2.2e-16
alternative hypothesis: two.sided
sample estimates:
    prop 1     prop 2     prop 3     prop 4     prop 5     prop 6
0.06081081 0.04545455 0.14640199 0.07843137 0.28901734 0.22775801
Pairwise comparisons using Pairwise comparison of proportions
data:  c(one_ctrl$count[k], one_spo$count[k], one_lmn$count[k], one_double$count[k],  out
of c(tot_ctrl$count[k], tot_spo$count[k], tot_lmn$count[k], tot_double$count[k],
one_triple$count[k], one_sun$count[k]) out of     tot_triple$count[k], tot_sun$count[k])
  1       2       3       4       5
2 0.66674 -       -       -       -
3 0.01564 0.00021 -       -       -
4 0.66674 0.21027 0.01741 -       -
5 1.2e-06 2.0e-10 0.00021 8.2e-08 -
6 5.6e-05 4.8e-08 0.01446 1.1e-05 0.21027
P value adjustment method: BH
[1] "this is for zone 2"
6-sample test for equality of proportions without continuity correction
data:  c(one_ctrl$count[k], one_spo$count[k], one_lmn$count[k], one_double$count[k],  out
of c(tot_ctrl$count[k], tot_spo$count[k], tot_lmn$count[k], tot_double$count[k],
one_triple$count[k], one_sun$count[k]) out of     tot_triple$count[k], tot_sun$count[k])
X-squared = 207.72, df = 5, p-value < 2.2e-16
alternative hypothesis: two.sided
sample estimates:
    prop 1     prop 2     prop 3     prop 4     prop 5     prop 6
0.20851064 0.04891304 0.30373832 0.12745098 0.30000000 0.49450549
Pairwise comparisons using Pairwise comparison of proportions
data:  c(one_ctrl$count[k], one_spo$count[k], one_lmn$count[k], one_double$count[k],  out
of c(tot_ctrl$count[k], tot_spo$count[k], tot_lmn$count[k], tot_double$count[k],
one_triple$count[k], one_sun$count[k]) out of     tot_triple$count[k], tot_sun$count[k])
  1       2       3       4       5
2 6.8e-09 -       -       -       -
3 0.01345 < 2e-16 -       -       -
4 0.01817 0.00061 7.9e-08 -       -
5 0.03296 2.5e-16 0.99158 2.4e-06 -
6 1.3e-10 < 2e-16 1.1e-06 < 2e-16 2.2e-05
P value adjustment method: BH
[1] "this is for zone 3"
6-sample test for equality of proportions without continuity correction
data:  c(one_ctrl$count[k], one_spo$count[k], one_lmn$count[k], one_double$count[k],  out
of c(tot_ctrl$count[k], tot_spo$count[k], tot_lmn$count[k], tot_double$count[k],
one_triple$count[k], one_sun$count[k]) out of     tot_triple$count[k], tot_sun$count[k])
X-squared = 910.61, df = 5, p-value < 2.2e-16
alternative hypothesis: two.sided
sample estimates:
    prop 1     prop 2     prop 3     prop 4     prop 5     prop 6
0.71897810 0.03389831 0.84318182 0.24800000 0.33070866 0.91772152
Pairwise comparisons using Pairwise comparison of proportions
data:  c(one_ctrl$count[k], one_spo$count[k], one_lmn$count[k], one_double$count[k],  out
of c(tot_ctrl$count[k], tot_spo$count[k], tot_lmn$count[k], tot_double$count[k],
one_triple$count[k], one_sun$count[k]) out of     tot_triple$count[k], tot_sun$count[k])
  1       2       3       4       5
2 < 2e-16 -       -       -       -
3 0.00011 < 2e-16 -       -       -
4 < 2e-16 6.2e-16 < 2e-16 -       -
5 < 2e-16 < 2e-16 < 2e-16 0.02978 -
6 5.7e-10 < 2e-16 0.00353 < 2e-16 < 2e-16
P value adjustment method: BH
[1] "this is for zone 4"
```

```
6-sample test for equality of proportions without continuity correction
data:  c(one_ctrl$count[k], one_spo$count[k], one_lmn$count[k], one_double$count[k],  out
of c(tot_ctrl$count[k], tot_spo$count[k], tot_lmn$count[k], tot_double$count[k],
one_triple$count[k], one_sun$count[k]) out of     tot_triple$count[k], tot_sun$count[k])
X-squared = 827.93, df = 5, p-value < 2.2e-16
alternative hypothesis: two.sided
sample estimates:
    prop 1     prop 2     prop 3     prop 4     prop 5     prop 6
0.56744186 0.06936416 0.89830508 0.21660650 0.36111111 1.00000000
Pairwise comparisons using Pairwise comparison of proportions
data:  c(one_ctrl$count[k], one_spo$count[k], one_lmn$count[k], one_double$count[k],  out
of c(tot_ctrl$count[k], tot_spo$count[k], tot_lmn$count[k], tot_double$count[k],
one_triple$count[k], one_sun$count[k]) out of     tot_triple$count[k], tot_sun$count[k])
  1       2       3       4       5
2 < 2e-16 -       -       -       -
3 < 2e-16 < 2e-16 -       -       -
4 4.1e-15 2.1e-07 < 2e-16 -       -
5 0.00077 8.9e-14 < 2e-16 0.00536 -
6 < 2e-16 < 2e-16 6.8e-07 < 2e-16 < 2e-16
P value adjustment method: BH
[1] "this is for zone 5"
6-sample test for equality of proportions without continuity correction
data:  c(one_ctrl$count[k], one_spo$count[k], one_lmn$count[k], one_double$count[k],  out
of c(tot_ctrl$count[k], tot_spo$count[k], tot_lmn$count[k], tot_double$count[k],
one_triple$count[k], one_sun$count[k]) out of     tot_triple$count[k], tot_sun$count[k])
X-squared = 327.26, df = 5, p-value < 2.2e-16
alternative hypothesis: two.sided
sample estimates:
    prop 1     prop 2     prop 3     prop 4     prop 5     prop 6
0.17391304 0.04705882 0.66666667 0.11206897 0.30379747 0.98809524
Pairwise comparisons using Pairwise comparison of proportions
data:  c(one_ctrl$count[k], one_spo$count[k], one_lmn$count[k], one_double$count[k],  out
of c(tot_ctrl$count[k], tot_spo$count[k], tot_lmn$count[k], tot_double$count[k],
one_triple$count[k], one_sun$count[k]) out of     tot_triple$count[k], tot_sun$count[k])
  1       2       3       4       5
2 0.0019  -       -       -       -
3 3.7e-13 < 2e-16 -       -       -
4 0.2813  0.0738  < 2e-16 -       -
5 0.0738  9.4e-08 4.2e-07 0.0019  -
6 < 2e-16 < 2e-16 5.0e-08 < 2e-16 < 2e-16
P value adjustment method: BH
```

**For comparing the percentage of X-chromosome pairing or complete synapsis; related to Supplementary Figure 6c:**

```
#Pairing:
> res1
2-sample test for equality of proportions with continuity correction
data:  c(9, 30) out of c(201, 175)
X-squared = 14.808, df = 1, p-value = 0.000119
alternative hypothesis: two.sided
95 percent confidence interval:
 -0.1947299 -0.0585750
sample estimates:
    prop 1     prop 2
0.04477612 0.17142857
> res2
2-sample test for equality of proportions with continuity correction
data:  c(150, 224) out of c(274, 248)
X-squared = 79.374, df = 1, p-value < 2.2e-16
alternative hypothesis: two.sided
95 percent confidence interval:
 -0.4291007 -0.2824604
sample estimates:
   prop 1    prop 2
0.5474453 0.9032258
> res3
2-sample test for equality of proportions with continuity correction
data:  c(297, 269) out of c(307, 273)
X-squared = 1.2829, df = 1, p-value = 0.2574
alternative hypothesis: two.sided
95 percent confidence interval:
 -0.045824466  0.009981916
sample estimates:
   prop 1    prop 2
0.9674267 0.9853480
> res4
2-sample test for equality of proportions with continuity correction
data:  c(258, 172) out of c(263, 172)
X-squared = 1.8463, df = 1, p-value = 0.1742
alternative hypothesis: two.sided
95 percent confidence interval:
 -0.040324288  0.002301475
sample estimates:
   prop 1    prop 2
0.9809886 1.0000000
> res5
2-sample test for equality of proportions with continuity correction
data:  c(132, 59) out of c(135, 63)
X-squared = 1.1058, df = 1, p-value = 0.293
alternative hypothesis: two.sided
95 percent confidence interval:
 -0.03551592  0.11805560
sample estimates:
   prop 1    prop 2
0.9777778 0.9365079
> res1$p.value
[1] 0.000119012
> res2$p.value
[1] 5.141022e-19
> res3$p.value
[1] 0.2573633
> res4$p.value
[1] 0.1742091
> res5$p.value
[1] 0.2930057

*** when p < 0.001
```

```
## synapsis:
2-sample test for equality of proportions with continuity correction
data:  c(0, 5) out of c(209, 192)
X-squared = 3.5994, df = 1, p-value = 0.0578
alternative hypothesis: two.sided
95 percent confidence interval:
 -0.053565110  0.001481777
sample estimates:
    prop 1     prop 2
0.00000000 0.02604167
> res7
2-sample test for equality of proportions with continuity correction
data:  c(52, 131) out of c(280, 238)
X-squared = 73.308, df = 1, p-value < 2.2e-16
alternative hypothesis: two.sided
95 percent confidence interval:
 -0.4464952 -0.2829166
sample estimates:
   prop 1    prop 2
0.1857143 0.5504202
> res8
2-sample test for equality of proportions with continuity correction
data:  c(254, 259) out of c(307, 286)
X-squared = 7.1089, df = 1, p-value = 0.00767
alternative hypothesis: two.sided
95 percent confidence interval:
 -0.13579083 -0.02067485
sample estimates:
   prop 1    prop 2
0.8273616 0.9055944
> res9
2-sample test for equality of proportions with continuity correction
data:  c(248, 153) out of c(261, 178)
X-squared = 9.8798, df = 1, p-value = 0.001671
alternative hypothesis: two.sided
95 percent confidence interval:
 0.02845383 0.15282819
sample estimates:
   prop 1    prop 2
0.9501916 0.8595506
> res10
2-sample test for equality of proportions with continuity correction
data:  c(61, 26) out of c(135, 64)
X-squared = 0.20501, df = 1, p-value = 0.6507
alternative hypothesis: two.sided
95 percent confidence interval:
 -0.1126318  0.2038355
sample estimates:
   prop 1    prop 2
0.4518519 0.4062500
> res6$p.value
[1] 0.05780018
> res7$p.value
[1] 1.109453e-17
> res8$p.value
[1] 0.007670096
> res9$p.value
[1] 0.001671028
> res10$p.value
[1] 0.6507078
```

**Statistics for comparing the extent of asymmetry, L4440/*samp-1(RNAi)* in EP, linear regression/ANOVA, "Y" being normalized intensity, "X" being angles from 1-180° (combined from 0-360°), "1" being L4440, "2" being samp-1 RNAi; related to Supplementary Figure 13b:**

```
> fit <- lm(Y ~ X * id,data=df_fit)
> anova(fit)
Analysis of Variance Table
Response: Y
            Df  Sum Sq Mean Sq F value    Pr(>F)    
X            1  46.980  46.980  936.15 < 2.2e-16 ***
id           1  11.414  11.414  227.45 < 2.2e-16 ***
X:id         1   9.651   9.651  192.31 < 2.2e-16 ***
Residuals 6280 315.161   0.050                      
---
Signif. codes:  0 '***' 0.001 '**' 0.01 '*' 0.05 '.' 0.1 ' ' 1
> print(fit)
Call:
lm(formula = Y ~ X * id, data = df_fit)
Coefficients:
(Intercept)            X           id         X:id  
  1.1572230    0.0005914    0.2182465   -0.0014932  
```

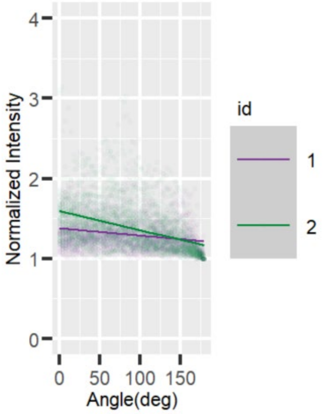

Statistics for comparing the extent of asymmetry, L4440/*lem-2(RNAi)* in EP, linear regression/ANOVA, “Y” being normalized intensity, “X” being angles from 1-180° (combined from 0-360°), “1” being L4440, “2” being lem-2 RNAi; related to Supplementary Figure 13d:

```
Analysis of Variance Table
Response: Y
            Df Sum Sq Mean Sq F value    Pr(>F)
X            1  67.01  67.006 1801.59 < 2.2e-16 ***
id           1  27.54  27.538  740.43 < 2.2e-16 ***
X:id         1  20.63  20.632  554.72 < 2.2e-16 ***
Residuals 9055 336.78   0.037
---
Signif. codes:  0 ‘***’ 0.001 ‘**’ 0.01 ‘*’ 0.05 ‘.’ 0.1 ‘ ’ 1
> print(fit)
Call:
lm(formula = Y ~ X * id, data = df_fit)
Coefficients:
(Intercept)            X           id         X:id
  1.0606633    0.0009627    0.2777642   -0.0018549

OR:
> res.aov3 <- aov(Y ~ X * id, data = df_fit)
> summary(res.aov3)
              Df Sum Sq Mean Sq F value Pr(>F)
X              1   67.0   67.01  1801.6 <2e-16 ***
id             1   27.5   27.54   740.4 <2e-16 ***
X:id           1   20.6   20.63   554.7 <2e-16 ***
Residuals   9055  336.8    0.04
---
Signif. codes:  0 ‘***’ 0.001 ‘**’ 0.01 ‘*’ 0.05 ‘.’ 0.1 ‘ ’ 1
```

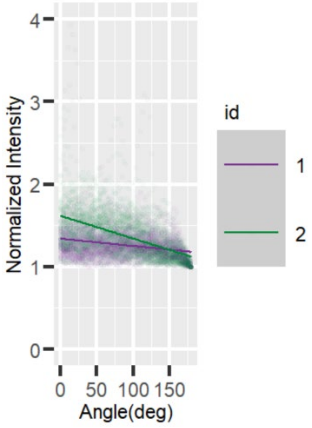
