## Supplementary Tables for "A cooperative network at the nuclear envelope counteracts LINC-mediated forces during oogenesis in *C. elegans*"

**Supplementary Table 1.** Quantification of brood size, embryo viability, and male self-progeny of worm strains with new AID alleles generated in this study.

| Strains | Auxin | n | Eggs laid (±SD) | Egg viability (±SD) %* | Male progeny (±SD) % |
|---|---|---|---|---|---|
| *$P_{sun-1}$::TIR1IV* | - | 6 | 225.50 ± 18.92 | 106.07 ± 3.22 | 0.00 ± 0.00 |
| *lmn-1::AID::V5 I;$P_{sun-1}$::TIR1 IV* | - | 7 | 252.14 ± 30.39 | 106.37 ± 5.07 | 0.09 ± 0.24 |
| *$P_{sun-1}$::TIR1* | + (from L1) | 4 | 228.00 ± 26.47 | 106.80 ± 3.84 | 0.00 ± 0.00 |
| *lmn-1::AID::V5 I;$P_{sun-1}$::TIR1 IV* | + (from L1) | 6 | 83.33 ± 45.78 | 9.60 ± 23.53 | 0.00 ± 0.00 |
| *$P_{sun-1}$::TIR1 IV* | + (from L4) | 3 | 245.67 ± 43.75 | 107.97 ± 2.34 | 0.12 ± 0.21 |
| *lmn-1::AID::V5 I;$P_{sun-1}$::TIR1 IV* | + (from L4) | 3 | 39.67 ± 44.00 | 0.00 ± 0.00 | 0.00 ± 0.00 |
| *HA::AID::zyg-12 II; $P_{sun-1}$::TIR1 IV* | - | 3 | 254.33 ± 76.42 | 104.10 ± 4.73 | 0.00 ± 0.00 |
| *sun-1::AID::V5 V; $P_{sun-1}$::TIR1 IV* | - | 6 | 225.50 ± 53.46 | 81.27 ± 21.71 | 0.71 ± 0.74 |
| *HA::AID::zyg-12 II; $P_{sun-1}$::TIR1 IV* | + (from L1) | 3 | 0.00 ± 0.00 | 0.00 ± 0.00 | 0.00 ± 0.00 |
| *sun-1::AID::V5 V; $P_{sun-1}$::TIR1 IV* | + (from L1) | 5 | 31.00 ± 26.46 | 0.00 ± 0.00 | 0.00 ± 0.00 |
| *HA::AID::zyg-12 II; $P_{sun-1}$::TIR1 IV* | + (from L4) | 3 | 0.00 ± 0.00 | 0.00 ± 0.00 | 0.00 ± 0.00 |
| *sun-1::AID::V5 V; $P_{sun-1}$::TIR1 IV* | + (from L4) | 3 | 33.33 ± 35.34 | 0.00 ± 0.00 | 0.00 ± 0.00 |
| *emr-1(gk119) I; lem-2::HA::AID II; $P_{sun-1}$::TIR1 IV* | - | 6 | 216.33 ± 22.47 | 106.42 ± 2.36 | 0.07 ± 0.17 |

* Egg viability of >100% is due to occasionally missed embryos when counting.

**Supplementary Table 2.** New alleles generated in this study.

| Allele | Genotype | Information about mutagenesis |
|---|---|---|
| ***ie137*** | *lmn-1(ie137[lmn-1::AID::V5])* | Internally tagged; generated using *dpy-10* Co-CRISPR in *ieSi38[sun-1p::TIR1::mRuby::sun-1 3'UTR, Cbr-unc-119(+)] IV* |
| ***ie138*** | *zyg-12(ie138[HA::AID::zyg-12])* | generated using *dpy-10* Co-CRISPR in *ieSi38[sun-1p::TIR1::mRuby::sun-1 3'UTR, Cbr-unc-119(+)] IV* |
| ***ie139*** | *sun-1(ie139[sun-1::AID::V5])* | Internally tagged; generated using *dpy-10* Co-CRISPR in *ieSi38[sun-1p::TIR1::mRuby::sun-1 3'UTR, Cbr-unc-119(+)] IV* |
| ***ie140*** | *lem-2(ie140[lem-2::HA::AID])* | generated using *dpy-10* Co-CRISPR in *ieSi38[sun-1p::TIR1::mRuby::sun-1 3'UTR, Cbr-unc-119(+)] IV* |

**Supplementary Table 3.** Genotypes of worm strains generated and used in this study.

| Strains | Source | Identifier |
|---|---|---|
| C.elegans: bcIs39(ced-1::GFP) V | Caenorhabditis Genetics Center | CA195 (MD701) |
| C.elegans: syp-1 (me17) bcIs39(ced-1::GFP) V/ nT1 [qIs51] (IV;V) | Bhalla et al., 2005 | CA885 |
| C.elegans: ieDf2/mIs11 IV | Harper et al., 2011; Caenorhabditis Genetics Center | CA998 |
| C. elegans: ieSi38[sun-1p::TIR1::mRuby::sun-1 3' UTR, Cbr-unc-119 (+)] IV | Zhang et al., 2015; Caenorhabditis Genetics Center | CA1199 |
| C.elegans: ieSi64[gld-1p::TIR1::mRuby::gld-1 3'UTR, Cbr-unc-119(+)] II; unc-119(ed3) III | Zhang et al., 2015; Caenorhabditis Genetics Center | CA1352 |
| C.elegans: ieSi65[sun-1p::TIR1::sun-1 3'UTR, Cbr-unc-119(+)] II; unc-119(ed3) III | Zhang et al., 2015; Caenorhabditis Genetics Center | CA1353 |
| C. elegans: meIs8[pie-1p::GFP::cosa-1, unc-119(+)] II; spo-11(ie59[spo-11::AID::3xFLAG]), ieSi38[sun-1p::TIR1::mRuby::sun-1 3'UTR, Cbr-unc-119(+)] IV | Zhang et al., 2018; Caenorhabditis Genetics Center | CA1423 |
| C. elegans: lmn-1(ie137[lmn-1::AID::V5]) I; unc-119 (ed3) III; ieSi38 [$P_{sun-1}$::TIR1::mRuby::sun-1 3'UTR, cb-unc-119(+)] IV | This paper | CA1532 |
| C. elegans: lmn-1(ie137[lmn-1::AID::V5]) I; unc-119 (ed3) III; ieSi38 [$P_{sun-1}$::TIR1::mRuby::sun-1 3'UTR, cb-unc-119(+)] IV; bcIs39 (ced-1::GFP) V | This paper | CA1561 |
| C. elegans: lmn-1(ie137[lmn-1::AID::V5]) I; ;ced-4(n1162) III; ieSi38 [$P_{sun-1}$::TIR1::mRuby::sun-1 3'UTR, cb-unc-119(+)] IV | This paper | CA1562 |
| C. elegans: lmn-1(ie137[lmn-1::AID::V5]) I; ieSi65[sun-1p::TIR1::sun-1 3'UTR, Cbr-unc-119(+)] II; unc-119(ed3) III; ieSi21 (sun-1::mRuby) IV | This paper | CA1563 |
| C. elegans: unc-119 (ed3) III; ieSi38 [$P_{sun-1}$::TIR1::mRuby::sun-1 3'UTR, cb-unc-119(+)] IV; sun-1(ie139[sun-1::AID::V5]) V | This paper | CA1564 |
| C. elegans: lmn-1(ie137[lmn-1::AID::V5]) I; ieSi64[gld-1p::TIR1::mRuby::gld-1 3'UTR, Cbr-unc-119(+)] II; unc-119(ed3) III; sun-1(ie139[sun-1::AID::V5]) V | This paper | CA1565 |
| C. elegans: zyg-12(ie138[HA::AID::zyg-12]) II; unc-119 (ed3) III; ieSi38 [$P_{sun-1}$::TIR1::mRuby::sun-1 3'UTR, cb-unc-119(+)] IV | This paper | CA1566 |
| C. elegans: lmn-1(ie137[lmn-1::AID::V5]) I; zyg-12(ie138[HA::AID::zyg-12]) II; unc-119 (ed3) III; ieSi38 [$P_{sun-1}$::TIR1::mRuby::sun-1 3'UTR, cb-unc-119(+)] IV | This paper | CA1567 |
| C. elegans: lmn-1(ie137[lmn-1::AID::V5]), emr-1(gk119) I; unc-119 (ed3) III; ieSi38 [$P_{sun-1}$::TIR1::mRuby::sun-1 3'UTR, cb-unc-119(+)] IV | This paper | CA1568 |
| C. elegans: lem-2(ie140[lem-2::HA::AID]) II; ieSi64[gld-1p::TIR1::mRuby::gld-1 3'UTR, Cbr-unc-119(+)] II; unc-119(ed3) III; | This paper | CA1569 |
| C. elegans: lmn-1(ie137[lmn-1::AID::V5]) I; lem-2(ie140[lem-2::HA::AID]), ieSi64[gld-1p::TIR1::mRuby::gld-1 3'UTR, Cbr-unc-119(+)] II; unc-119(ed3) III; | This paper | CA1570 |
| C. elegans: lmn-1(ie137[lmn-1::AID::V5]), emr-1(gk119) I; unc-119 (ed3) III; ieSi38 [$P_{sun-1}$::TIR1::mRuby::sun-1 3'UTR, cb-unc-119(+)] IV; sun-1(ie139[sun-1::AID::V5]) V | This paper | CA1571 |
| C. elegans: syp-3 (ok857) I; ieSi64[gld-1p::TIR1::mRuby::gld-1 3'UTR, Cbr-unc-119(+)] II; unc-119(ed3) III; ieSi19 (mRuby::SYP-3),ojIs9 [zyg-12(all)::GFP + unc-119(+)] IV; sun-1(ie139[sun-1::AID::V5]) V | This paper | CA1572 |
| C. elegans: lmn-1(ie137[lmn-1::AID::V5]) I; ieSi64[gld-1p::TIR1::mRuby::gld-1 3'UTR, Cbr-unc-119(+)] II; unc-119(ed3) III; ieSi19 (mRuby::SYP-3),ojIs9 [zyg-12(all)::GFP + unc-119(+)] IV; sun-1(ie139[sun-1::AID::V5]) V | This paper | CA1573 |
| C. elegans: lmn-1(ie137[lmn-1::AID::V5]) I; ieSi64[gld-1p::TIR1::mRuby::gld-1 3'UTR, Cbr-unc-119(+)] II; unc-119(ed3) III; ieSi19 (mRuby::SYP-3),ojIs9 [zyg-12(all)::GFP + unc-119(+)] IV; | This paper | CA1574 |
| C. elegans: lmn-1(ie137[lmn-1::AID::V5]), emr-1(gk119) I; ieSi64[gld-1p::TIR1::mRuby::gld-1 3'UTR, Cbr-unc-119(+)] II; unc-119(ed3) III; ieSi19 (mRuby::SYP-3),ojIs9 [zyg-12(all)::GFP + unc-119(+)] IV; | This paper | CA1575 |
| C. elegans: lmn-1(ie137[lmn-1::AID::V5]) I; meIs8[pie-1p::GFP::cosa-1, unc-119(+)] II; unc-119 (ed3) III; ieSi38 [$P_{sun-1}$::TIR1::mRuby::sun-1 3'UTR, cb-unc-119(+)] IV | This paper | CA1576 |
| C. elegans: lmn-1(ie137[lmn-1::AID::V5]) I; meIs8[pie-1p::GFP::cosa-1, unc-119(+)] II; unc-119 (ed3) III; spo-11(ie59[spo-11::AID::3xFLAG]), ieSi38 [$P_{sun-1}$::TIR1::mRuby::sun-1 3'UTR, cb-unc-119(+)] IV | This paper | CA1577 |
| C. elegans: lmn-1(ie137[lmn-1::AID::V5]) I; unc-119 (ed3) III; spo-11(ie59[spo-11::AID::3xFLAG]), ieSi38 [$P_{sun-1}$::TIR1::mRuby::sun-1 3'UTR, cb-unc-119(+)] IV; sun-1(ie139[sun-1::AID::V5]) V | This paper | CA1578 |
| C. elegans: zyg-12(ie138[HA::AID::zyg-12]) II; unc-119 (ed3) III; ieSi38 [$P_{sun-1}$::TIR1::mRuby::sun-1 3'UTR, cb-unc-119(+)] IV; bcIs39 (ced-1::GFP) V | This paper | CA1579 |

**Supplementary Table 4.** Sequences of gRNAs, repair templates, and DNA primers used to genotype edited progeny.

| Transgenes | gRNAs and repair templates (mostly gBlock) | Genotyping primer names | Primer sequences | Fragment sizes |
|---|---|---|---|---|
| *lmn-1::AID::V5* in *ie137* | 5’ – AGAAGTTCGTCACAAGAGAC – 3’; 5’ - TCTGGAAGAAGATCTCGCTTTTGCTCTTCAACAGCACAAGGGAGAACTTGAAGAAGTTCGcCACAAGAGgCAGGTCGACATGACAACCTACGGCGGCGGAGGATCCatgcctaaagatccagccaaacctccggccaaggcacaagttgtgggatggccaccggtgagatcataccggaagaacgtgatggtttcctgccaaaaatcaagcggtggcccggaggcggcggcgttcgtgaagggaggatccggaGGAAAGCCAATTCCAAACCCACTTCTTGGACTCGACTCCACCGCCAAGCAGATTAATGATGAGTATCAATCTAAGCTT -3’ | oCL41 (F) | 5' - AAA GCA GAA CAT CAC TCT TCG TGA CAC CGT AGA AG -3’; | WT, 277bp; inserted, 481bp |
| | | oCL42 (R) | 5' - TTT GAT GCA AAT TGT TCT TGA ACT GAG CAC GCA TCT C -3’ | |
| *HA::AID::zyg-12* in *ie138* | 5’ – GAATCTGAGTCGTCAGACAA – 3’; 5’ – aaaatctatcaatttctttttttcagaacaaaatcatgTACCCATACGATGTTCCAGATTACGCTggaggatccggaatgcctaaagatccagccaaacctccggccaaggcacaagttgtgggatggccaccggtgagatcataccggaagaacgtgatggtttcctgccaaaaatcaagcggtggcccggaggcggcggcgttcgtgaagGGCGGCGGAGGATCCGGAGGAGGAGGCAGTGGAGGCGGCGGTTCTGGCGGTGGCGGCTCAGGCGGAGGTGGATCGTTAGACCTGACAAACAAAGAGTCCGAGTCTTCAGACAACGGAAATAGCAAGTACGAAGATTCCATAGACGGACGA – 3’ | oCL95 (F) | 5’ – TTGTAAACTCTACCAGCCT -3’; | WT, 401bp; inserted, 650bp |
| | | oCL96 (R) | 5’ – TCAGAGGTAGTTTAGTGGC -3’; | |
| *sun-1::AID::V5* in *ie139* | 5’ – GCTGGAATATCGCATTCGCA – 3’; 5’ – TACAAGGAGCATTTTAGCTACAAAGAAATCACTTCGATGAAGAAGGAAATGTGGTATGACTGGCTGGAATATCGCATcCGtGGCGGCGGAGGATCCatgcctaaagatccagccaaacctccggccaaggcacaagttgtgggatggccaccggtgagatcataccggaagaacgtgatggtttcctgccaaaaatcaagcggtggcccggaggcggcggcgttcgtgaagggaggatccggaGGAAAGCCAATTCCAAACCCACTTCTTGGACTCGACTCCACCATGGTTCGGCGTCGTTTTGTTCCAACGTGGGCCCAGTTTAAACGTACTCTT – 3’ | oCL101 (F) | 5'- CTT CGA TGA AGA AGG AAA TGT GGT ATG ACT GGC -3'; | WT, 456bp; inserted, 660bp |
| | | oCL102 (R) | 5'- CTC TTC GAT TGC CGA CTC TTT CCA TCC TTT -3'; | |
| *lem-2::HA::AID* in *ie140* | 5’ – TGTGCCGTGTGGAAGTGGAT – 3’; 5’ – CTACCGATGTTCTTGTGCTTCCGTCTGGAAATGAGTGcGCtGTcTGGAAaTGGATCGGAAATCAGTCTCAGAAGAGATGGTACCCATACGATGTTCCAGATTACGCTggaggatccggaatgcctaaagatccagccaaacctccggccaaggcacaagttgtgggatggccaccggtgagatcataccggaagaacgtgatggtttcctgccaaaaatcaagcggtggcccggaggcggcggcgttcgtgaagTAGatcattgttttgctgtataatttttcgatttt– 3’ | oCL137 (F) | 5’ – GAAGCTCTACGAGCTCATC – 3’; | WT, 379bp; inserted, 553bp |
| | | oCL138 (R) | 5’ – gtcattgtgataccttaggc – 3’; | |
| *dpy-10* | 5’ – GCTACCATAGGCACCACGAG – 3’; 5’ – ATACGGCAAGATGAGAATGACTGGAAACCGTACCGCATGCGGTGCCTATGGTAGCGGAGCTTCACATGGCTTCAGA – 3’ (ssDNA repair template) | N.A. | N.A. | N.A. |

**Supplementary Table 5.** RNAi clones (Ahringer library) used in this study.

| Target gene | GenePairs Name | Plate | Well |
|---|---|---|---|
| *dnc-1* | ZK593.5 | 111 | B12 |
| *dlc-1* | T26A5.9 | 75 | H6 |
| *lis-1* | T03F6.5 | 88 | F4 |
| *dhc-1* | T21E12.4 | 4 | H2 |
| *dli-1* | C39E9.14 | 116 | A3 |
| *dylt-1* | F13G3.4 | 11 | E6 |
| *lem-2* | W01G7.5 | 63 | B2 |
| *samp-1* | T24F1.2 | 58 | A6 |
| *lmn-1* | DY3.2 | 14 | D12 |
